# Rapamycin Mitigates a Sex-biased Convergent Aging Trajectory

**DOI:** 10.64898/2026.07.02.736117

**Authors:** Tzu-Chiao Lu, Chung-Yi Liang, Ye-Jin Park, Niccole Auld, Tyler Jackson, Zhiyong Yin, Erin Harrison, Bo Sun, Mujeeb Qadiri, Norbert Perrimon, Ao-Lin Hsu, Yanyan Qi, Hongjie Li

**Affiliations:** Department of Molecular and Human Genetics, Baylor College of Medicine, Houston, TX, USA; Institute of Biochemistry and Molecular Biology, National Yang Ming Chiao Tung University, Taipei, Taiwan; Development, Disease Models & Therapeutics Graduate Program, Baylor College of Medicine, Houston, USA; Cancer and Cell Biology Graduate Program, Baylor College of Medicine, Houston, USA; Department of Genetics, Blavatnik Institute, Harvard Medical School, Harvard University, Boston, MA, USA; Howard Hughes Medical Institute, Boston, USA; Institute of Molecular Biology, Academia Sinica, Taipei, Taiwan

**Keywords:** aging, longevity, Rapamycin, fly cell atlas, *Drosophila*, Convergent Aging Trajectory

## Abstract

Rapamycin extends lifespan across species, yet its cell-type-specific benefits and vulnerabilities remain unclear at whole-organism scale. Here, we present the Rapamycin Fly Cell Atlas (Rapa-FCA), a whole-organism single-nucleus transcriptomic atlas of *Drosophila* spanning both sexes, multiple ages, 18 cell classes, and 181 cell types. Rapamycin elicited a highly heterogeneous response, with prominent effects in reproductive, digestive, and neuromuscular systems and modest responses in most neuronal populations. Across diverse tissues, we identified a rapamycin-sensitive Convergent Aging Trajectory (CAT), marked by *Fkbp12* enrichment and mTORC1-linked metabolic programs, including glycolysis and lipid synthesis. CAT-high nuclei accumulated with age and were preferentially reduced by rapamycin, especially in females, consistent with stronger female lifespan extension. By integrating CAT abundance, aging-clock predictions, and nucleus-ratio changes, we mapped sex- and cell-type-specific geroprotection effects of rapamycin. Together, the Rapa-FCA provides an organism-wide framework for resolving how rapamycin reshapes cellular aging across sex, tissue, and cellular state.

## Introduction

Aging is a complex biological process characterized by gradual decline and dysfunction of systems and organs, thereby increasing disease vulnerability and mortality. As the global population ages, interventions that not only extend lifespan but also improve healthspan have become a central focus of biomedical research. Among all known interventions, rapamycin stands out as one of the most promising interventions (Jiang et al., 2025). As an inhibitor of the mechanistic target of rapamycin (mTOR) pathway, rapamycin has been shown to extend lifespan across species ranging from yeast and worms to flies and mammals (Bjedov et al., 2010; Chen et al., 2013; Harrison et al., 2009; Kennedy and Lamming, 2016; Powers et al., 2006). Its ability to modulate a central nutrient-sensing pathway has fueled excitement for its potential translation to humans.

The conserved lifespan-extending effects of rapamycin suggest that suppression of mTOR signaling is a central mechanism for aging intervention. However, aging-associated mTOR activity is unlikely to increase uniformly across the organism. Instead, increased or dysregulated activity of mTOR-associated pathways, especially those controlled by mTOR complex 1 (mTORC1), has been observed in selected tissues and cellular contexts across flies, mice, rats, and humans (Calhoun et al., 2016; Chen et al., 2009; Choy et al., 2025; Joseph et al., 2019; Leontieva et al., 2014; Portier et al., 2024; Tang et al., 2019; Yang et al., 2012). Given the importance of mTOR in diverse age-related metabolic disorders and diseases (Saxton and Sabatini, 2017), defining where and how mTOR activity changes during aging may inform strategies to modulate aging itself and identify therapeutic opportunities for age-associated disorders.

Despite its robust lifespan-extending effects, rapamycin does not act uniformly across biological contexts. Its impact varies with dose, treatment timing, sex, tissue, and cellular state (Bjedov et al., 2010; Dou et al., 2017; Fan et al., 2015; Jiang et al., 2025; Mannick et al., 2018; Wilkinson et al., 2012). Also, chronic treatment has been associated with metabolic and reproductive trade-offs in some settings (Lamming et al., 2012; Liu et al., 2017; Zhu et al., 2019). These context-dependent effects raise a central unresolved question: how are rapamycin’s organismal benefits and vulnerabilities distributed across cell types? More specifically, it remains unclear which cell types mediate its geroprotective effects, which show neutral or potentially adverse responses, whether these responses reflect direct mTOR inhibition or indirect systemic remodeling, how sex shapes rapamycin efficacy, and what molecular features define rapamycin-sensitive cells. Addressing these questions requires a systematic, whole-organism view of rapamycin action at cellular resolution.

The fruit fly, *Drosophila melanogaster*, offers a powerful experimental model to address this challenge. Core nutrient-sensing pathways, including mTOR signaling, are highly conserved from flies to mammals, and some key components of TOR-associated signaling were first characterized in *Drosophila* (Montagne et al., 1999; Oldham et al., 2000; Saucedo et al., 2003; Saxton and Sabatini, 2017). With advances in single-cell RNA sequencing, flies now provide unparalleled resolution for mapping cell type-specific changes during aging across the whole fly. Our whole-organism single-nucleus RNA sequencing (snRNA-seq) platform enabled the construction of the Fly Cell Atlas (FCA), which cataloged over 250 cell types in the adult fruit fly (Li et al., 2022). Building on this foundation, we developed the Aging Fly Cell Atlas (AFCA), which identified cell-type–specific features of aging across the organism (Lu et al., 2023).

Here, we apply this platform to rapamycin intervention. We generated the Rapamycin Fly Cell Atlas (Rapa-FCA) using whole-organism snRNA-seq of male and female flies treated with rapamycin across aging. In total, we analyzed more than 505,000 high-quality nuclei across four ages, encompassing 18 broad cell classes and over 180 cell types. Our analysis reveals that reproductive, digestive, and neuromuscular systems are among the most responsive to rapamycin treatment, whereas most neuronal and glial cell types exhibit relatively modest changes. Importantly, rapamycin exerts profoundly sexually dimorphic effects, a divergence potentially rooted in the differential baseline expression of *Fkbp12*, the intracellular cofactor for rapamycin-mediated mTORC1 suppression. Most notably, we discovered that aging nuclei from diverse developmental origins converge toward a shared transcriptomic state enriched for mTORC1 signaling and *Fkbp12* expression, which we define as the Convergent Aging Trajectory (CAT). By integrating this convergent response with changes in nucleus ratio and cell type-specific aging clock predictions, we quantified rapamycin-associated geroprotection across cell types. This analysis revealed that many male cell types show neutral or negative responses early in life but gain protective effects later, whereas most female cell types benefit more consistently across early and late stages.

Together, these findings provide the first organism-wide, single-cell resolution map of rapamycin’s actions. The Rapa-FCA illuminates tissue-specific benefits and vulnerabilities, highlights sex-dependent responses, and establishes a new framework to refine rapamycin-based interventions for longevity. All data can be accessed, downloaded, and analyzed through https://hongjielilab.org/rapa-fca/.

## Results

### Constructing Rapamycin Fly Cell Atlas (Rapa-FCA)

To establish an organism-wide single-nucleus atlas for assessing rapamycin-mediated aging intervention, we first determined a rapamycin dose that reproducibly extends lifespan under our laboratory conditions. Previous *Drosophila* studies have used different rapamycin concentrations and reported dose-, sex-, and context-dependent effects on lifespan (Bjedov et al., 2010; Schinaman et al., 2019). Under our experimental conditions, the lower dose (20 μM) yielded a more robust lifespan extension for both sexes than a higher dose (200 μM). Consequently, we selected the 20 μM rapamycin for all downstream Rapa-FCA profiling. Consistent with previous reports (Bjedov et al., 2010; Schinaman et al., 2019), this treatment extended lifespan across both sexes, yielding a more pronounced geroprotective benefit in females (∼12% extension) than in males (∼7%) (**Figure 1a**).

**Figure 1.**
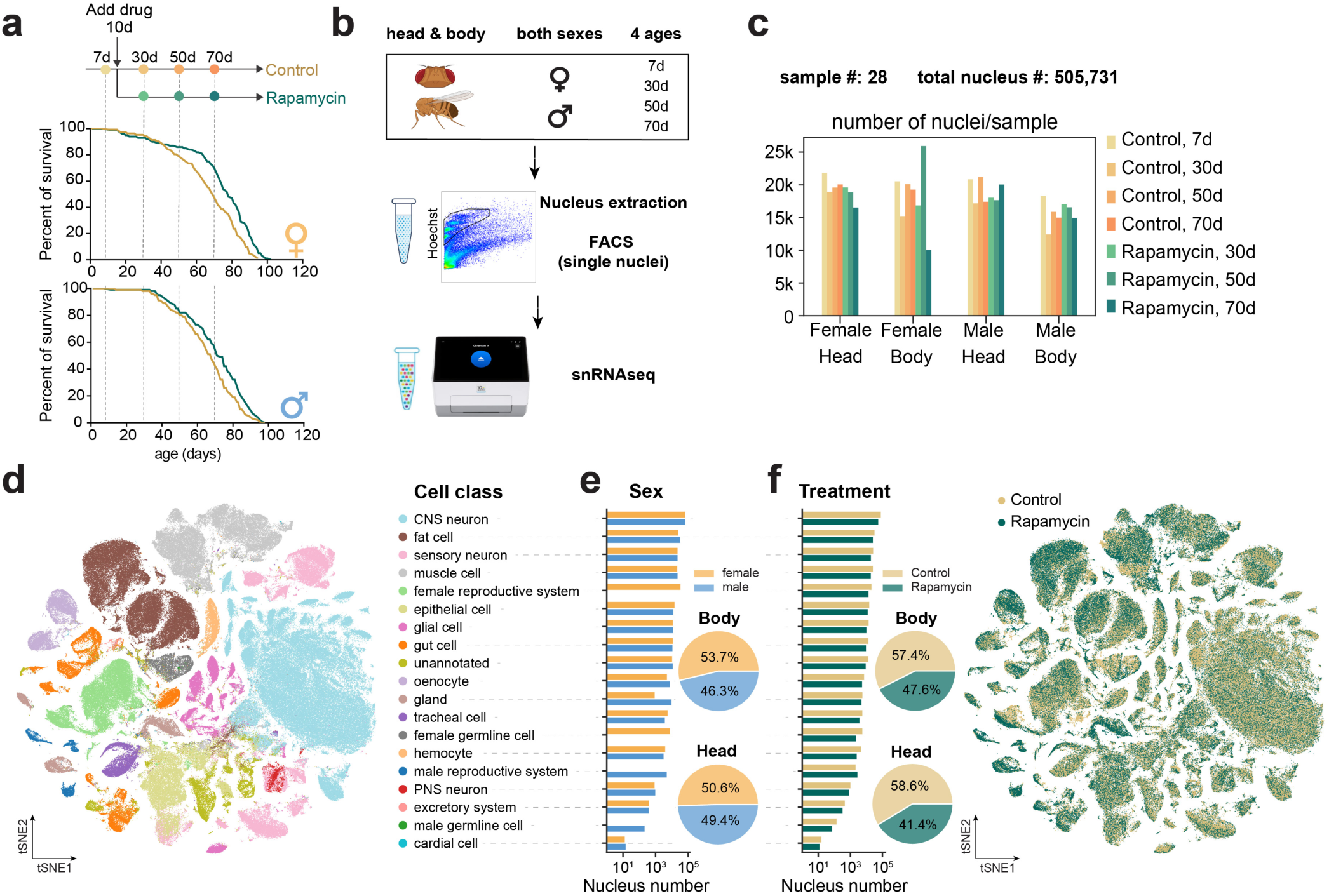
Overview of Rapa-FCA **a.** Lifespan analysis of female and male flies maintained on control or rapamycin-supplemented food. Rapamycin treatment was initiated at day 10, and flies were sampled at day 7, day 30, day 50, and day 70 for atlas construction. Survival curves are presented separately for females and males. Rapamycin treatment significantly extended median lifespan in both females (68.9 vs. 77.7 days; *p* = 2.1 × 10⁻⁷; n = 410 control, 409 rapamycin) and males (67.5 vs. 71.9 days; *p* = 0.0002; n = 311 control, 326 rapamycin). Statistical significance was determined using the log-rank (Mantel-Cox) test. **b.** Experimental design and workflow for whole-organism snRNA-seq profiling. Heads and bodies were collected separately from both sexes across age and treatment conditions. Hoechst-positive nuclei were isolated by FACS and captured using the 10x Genomics high-throughput 3′ gene-expression platform. **c.** Summary of the number of libraries and high-quality nuclei recovered across Rapa-FCA samples. **d.** t-distributed stochastic neighbor embedding (tSNE) visualization of Rapa-FCA nuclei, colored by 18 broad cell classes. **e.** Recovered nucleus numbers for each broad cell class, separated by sex. Pie charts show the overall proportions of nuclei recovered from females and males in body and head samples. **f.** Recovered nucleus numbers for each broad cell class, separated by treatment. Pie charts show the overall proportions of nuclei recovered from control and rapamycin-treated samples in body and head datasets. The tSNE visualization shows the distribution of control and rapamycin-treated nuclei across the integrated atlas.

We next generated whole-organism snRNA-seq profiles across sex, age, tissue compartment, and treatment. Young flies (day 7) were collected as pretreatment controls, whereas three later ages (day 30, day 50, and day 70) were collected from both control- and rapamycin-treated groups. Heads and bodies were processed separately for each sex and condition. After fluorescence-activated cell sorting (FACS)-assisted isolation of Hoechst-positive nuclei, we generated droplet-based snRNA-seq libraries using the 10x Genomics high-throughput platform (**Figure 1b**). In total, Rapa-FCA comprised 28 libraries and 505,731 high-quality nuclei after quality control (**Figure 1c** and **Extended Data Fig. 1**). We annotated cell identities by transferring labels from our previous atlas datasets (Li et al., 2022; Lu et al., 2023; Park et al., 2025), followed by manual validation using established cell-type-specific markers. This analysis identified 18 broad cell classes and 181 detailed cell types across the adult fly (**Figure 1d** and **Extended Data Fig. 2–3**). Major cell classes were broadly represented in both sexes, with expected differences in sex-specific reproductive cell types (**Figure 1e**). Across control and rapamycin-treated samples, the recovered nucleus composition was largely comparable, allowing systematic comparisons of transcriptomic responses across ages, sexes, and treatments (**Figure 1f**).

### Heterogeneous responses to rapamycin across the whole organism

To define how rapamycin reshapes gene expression across the adult fly, we compared rapamycin-treated flies with age- and sex-matched controls for each annotated cell type and ranked cell types by the number of differentially expressed genes (DEGs) (**Figure 2a**). Rapamycin-induced transcriptional changes were highly heterogeneous across the organism. Cell types with the largest DEG numbers were enriched in peripheral tissues, whereas most neuronal and glial populations showed comparatively modest responses. This pattern suggests that rapamycin, at current concentration, acts more strongly on peripheral tissues (see Discussion). Among the most responsive cell types, reproductive tissues were prominently represented, accounting for 7 of the top 15 affected cell types (**Figure 2a**) and including both female- and male-specific populations (**Figure 2b**). Several metabolically active or contractile somatic cell types, including the fat body, anterior enterocytes, oenocytes, ventral nervous cord (VNC), and indirect flight muscle (IFM), also showed high DEG numbers in both sexes.

**Figure 2.**
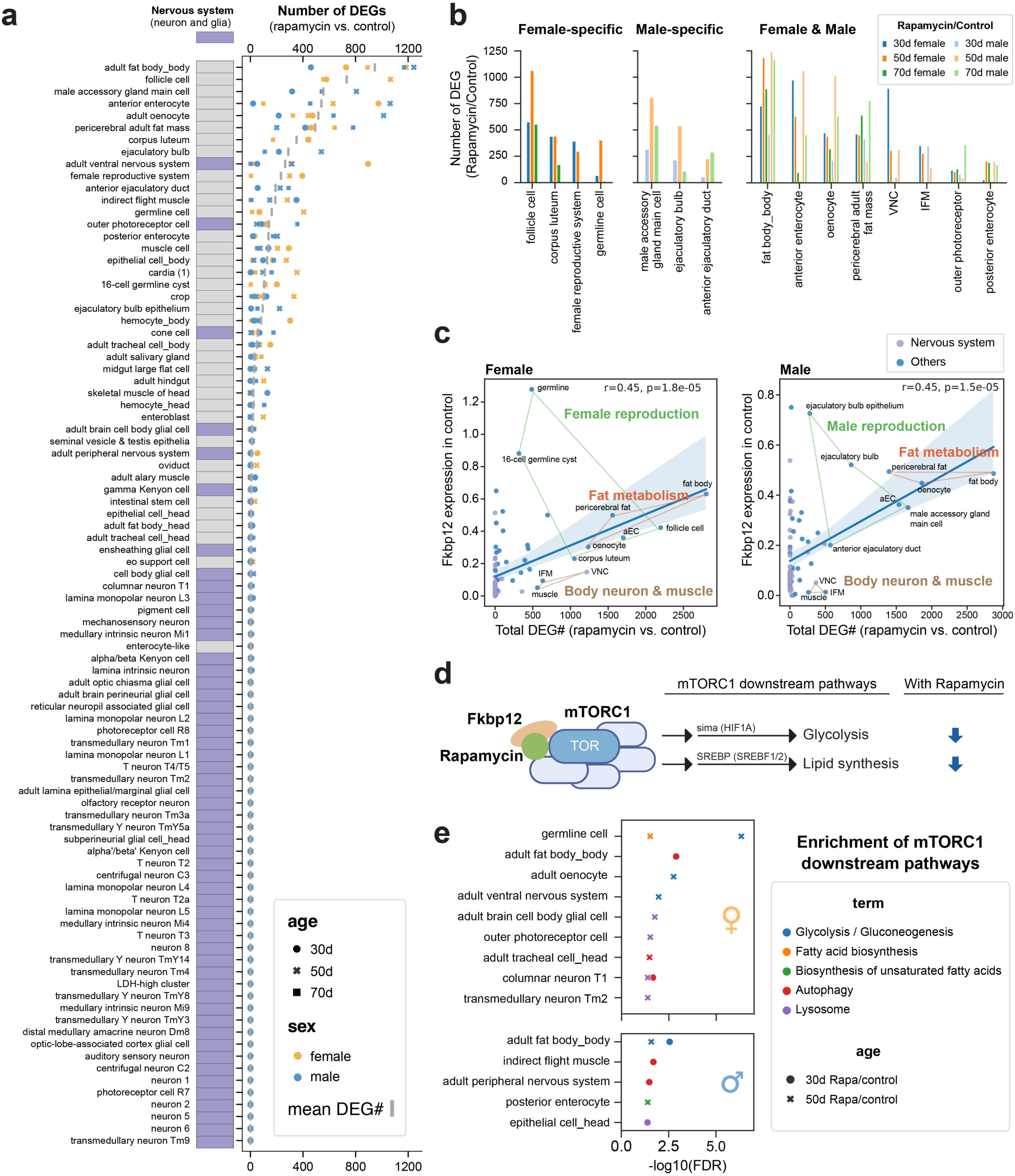
Cell-type- and sex-specific transcriptomic responses to rapamycin **a.** Ranking of rapamycin-responsive cell types based on the number of DEGs between rapamycin-treated and age- and sex-matched control samples. Cell types are ordered by mean DEG number across ages and sexes, with neuronal and glial populations indicated separately. **b.** Summary of highly responsive cell types classified as female-specific, male-specific, or shared between sexes. **c.** Relationship between basal *Fkbp12* expression in control samples and the magnitude of rapamycin-induced transcriptomic response across cell types. The correlation between the total number of DEGs and *Fkbp12* expression was assessed using the Pearson correlation coefficient. **d.** Schematic of rapamycin-mediated mTORC1 inhibition through Fkbp12 and selected downstream transcriptional programs. **e.** KEGG pathway enrichment analysis of rapamycin-induced DEGs across selected cell types.

Because Fkbp12 is the binding partner required for rapamycin-mediated inhibition of mTORC1 (Heitman et al., 1991; Sabatini et al., 1994), we hypothesized that variation in *Fkbp12* expression may contribute to cell-type-specific differences in rapamycin responsiveness. To test this possibility, we compared basal *Fkbp12* expression in control samples with the number of rapamycin-induced DEGs across cell types (**Figure 2c**). *Fkbp12* expression was positively correlated with DEG number, and this association was stronger than expected for randomly selected genes or most other FKBP family members (**Extended Data Fig. 4**). These results suggest that *Fkbp12* expression is linked to the magnitude of rapamycin-associated transcriptomic remodeling across cell types. Cell types with both high *Fkbp12* expression and high DEG numbers were enriched for reproductive and fat-associated tissues. In contrast, VNC and muscle cells exhibited high DEG numbers despite relatively low average *Fkbp12* expression, raising the possibility that rapamycin responsiveness in these tissues may arise from *Fkbp12*-independent functions or complex regulations.

We then examined whether rapamycin-responsive cell types showed evidence of altered mTORC1-associated transcriptional programs. mTORC1 regulates several downstream metabolic programs, including glycolysis and lipid biosynthesis, through transcriptional regulators such as sima/HIF1A and SREBP/SREBF1/2 (**Figure 2d**) (Dekanty et al., 2005; Liu and Sabatini, 2020; Porstmann et al., 2008). We therefore performed the Kyoto Encyclopedia of Genes and Genomes (KEGG) pathway enrichment analysis using cell-type- and sex-specific DEGs. Several responsive cell types were enriched for mTORC1-associated pathways, with glycolysis/gluconeogenesis showing the strongest enrichment in both females and males (**Figure 2e**). These pathway-enriched cell types partially overlapped with those showing high *Fkbp12* expression and high DEG numbers, supporting the idea that rapamycin-sensitive cell types are not randomly distributed across the organism but are associated with specific molecular and metabolic features. Together, these analyses identify reproductive, metabolic, and neuromuscular cell types as major sites of rapamycin-associated transcriptional remodeling and provide a framework for subsequent analyses of mTORC1-linked aging signatures.

### Rapamycin attenuates aging-associated shifts in reproductive cells

To determine whether rapamycin mitigates cell-type-specific aging, we first focused on the female germline, which showed high *Fkbp12* expression and enrichment of mTORC1-associated transcriptional programs in the initial analysis (**Figures 2c–e**). The *Drosophila* ovary contains multiple germline and somatic cell types that support oogenesis across progressive differentiation stages (**Figure 3a**). We therefore sub-selected germline nuclei and reconstructed their aging trajectories. In control flies, germline nuclei formed an age-associated lineage that was largely distinct from young nuclei and was enriched at day 50 (**Figure 3b** and **Extended Data Fig. 5a**). Strikingly, rapamycin treatment significantly decreased this aged lineage, demonstrating that mTOR inhibition suppresses the accumulation of this aged germline state. We defined the aged lineage based on age distribution and Leiden clustering in control samples and then quantified its abundance across ages and treatments. Using this approach, we observed an approximately 35% reduction in the aged germline lineage in rapamycin-treated flies compared with age-matched controls at day 50 (**Figure 3c**).

**Figure 3.**
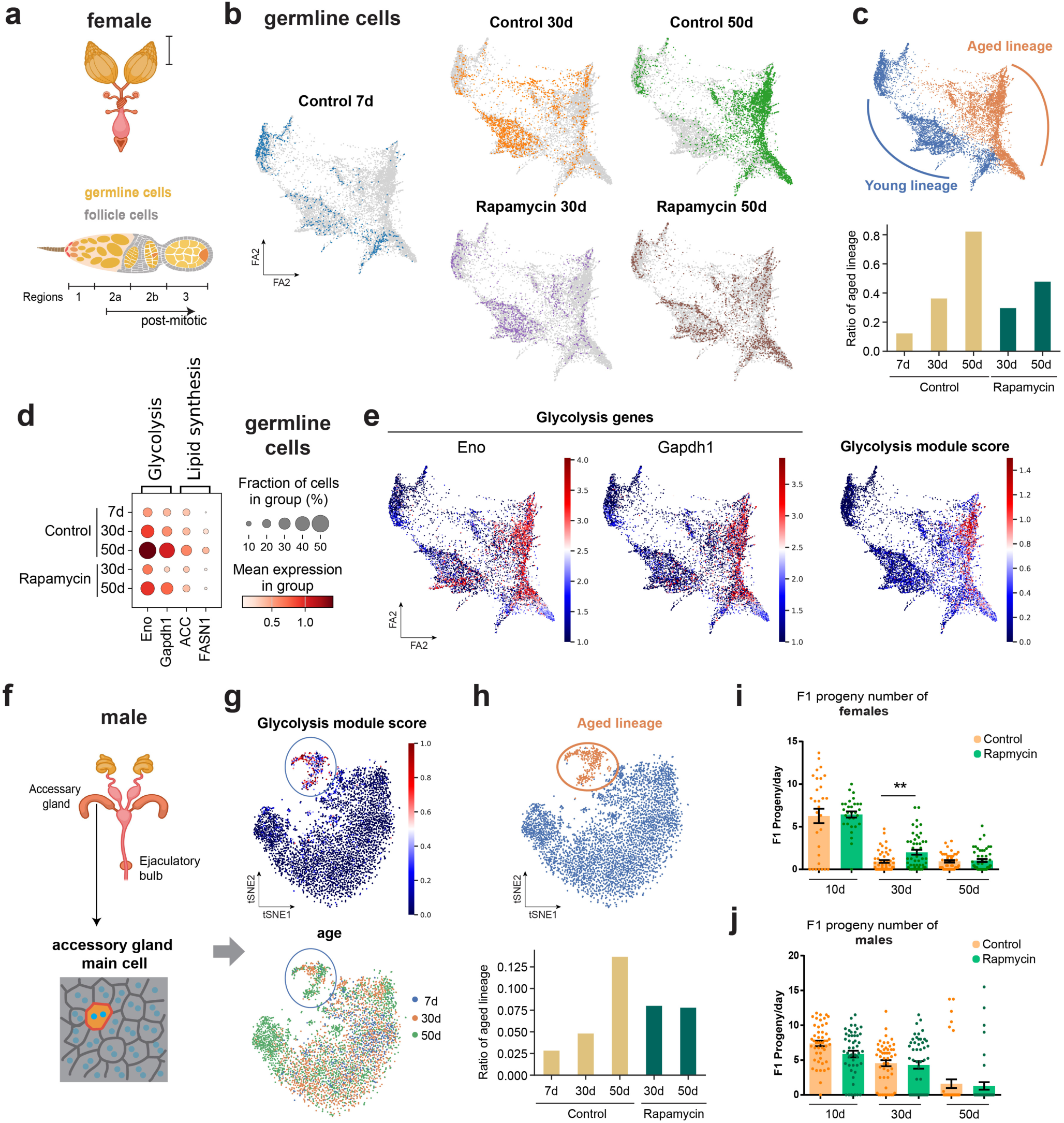
Rapamycin attenuates mTORC1-linked programs in reproductive cells **a.** Schematic of the female reproductive system and major ovary-associated cell populations, including germline and follicle-cell populations across oogenesis stages. **b.** Trajectory plot of female germline nuclei across age and treatment conditions. **c.** Definition and quantification of young and aged germline lineages. **d.** Expression of glycolysis and lipid-synthesis genes in female germline nuclei across age and treatment conditions. **e.** Trajectory plot of glycolysis gene expression and glycolysis module scores in female germline nuclei. **f.** Schematic of the male reproductive system, highlighting the accessory gland and ejaculatory bulb. **g.** tSNE visualization of male accessory gland main cells colored by glycolysis module score and age, showing the emergence of a glycolysis-high aged population. **h.** Definition and quantification of the aged lineage in male accessory gland main cells across age and treatment conditions. **i.** Female reproductive output measured by F1 progeny number per day across ages and treatments. Bars indicate mean ± SEM; **: unpaired t-test, p = 0.0019. **j.** Male reproductive output measured by F1 progeny number per day across ages and treatments.

Because this aged germline lineage was reduced by rapamycin, we next examined whether it was associated with mTORC1-linked transcriptional programs. Glycolysis and lipid-synthesis genes increased progressively with age in control germline nuclei, consistent with aging-associated activation of mTORC1-linked programs (**Figure 3d**) (Chen et al., 2009; Choy et al., 2025). Rapamycin treatment significantly reduced the expression of these genes at day 50, supporting attenuation of these pathways under rapamycin administration. Consistent with this interpretation, glycolysis and lipid-synthesis genes were enriched in the aged germline lineage compared with the non-aged population (**Figures 3e** and **Extended Data Fig. 5b**). Thus, aging in the female germline is accompanied by the emergence of a metabolically active aged lineage marked by mTORC1-linked transcriptional features, and this lineage is diminished by rapamycin treatment.

We next asked whether a similar aging-associated trajectory could be detected in male reproductive cells. We focused on male accessory gland main cells, which also showed high *Fkbp12* expression and high DEG numbers in response to rapamycin (**Figures 2c** and **3f**). Similar to the female germline, male accessory gland main cells contained an aged population that was transcriptionally separated from young nuclei and showed elevated glycolysis gene expression (**Figure 3g**). The abundance of this aged population increased with age in controls and was reduced by rapamycin at day 50 (**Figure 3h**). These findings suggest that reproductive cell types in both sexes can develop aging-associated transcriptional states enriched for mTORC1-linked metabolic programs, although the magnitude and timing of rapamycin-mediated mitigation differ between sexes.

Given the rapamycin-associated reduction of aged reproductive lineages, we next tested whether rapamycin treatment preserved reproductive output during aging. Rapamycin-treated females produced significantly more F1 progeny than age-matched controls, indicating improved reproductive capability in aged females (**Figure 3i**). In contrast, rapamycin did not significantly improve male reproductive output (**Figure 3j**). Notably, the aged-lineage ratio in male accessory gland main cells was slightly higher in rapamycin-treated flies than in controls at day 30, whereas aged-lineage ratios in the female germline were consistently lower after rapamycin treatment across ages (**Figures 3c** and **3h**). Together, these results indicate that rapamycin attenuates aging-associated, mTORC1-linked transcriptional states in reproductive cells, with a more consistent protective effect in the female germline that is accompanied by preserved female reproductive output.

### Sex-specific effects of rapamycin in VNC and muscle cells

We next examined cell types with high rapamycin-induced DEG numbers but relatively low *Fkbp12* expression, including VNC and muscle cells (**Figure 2c**). We reasoned that the strong transcriptomic responses in these cell types could arise either from small *Fkbp12*-high subpopulations or from indirect systemic effects of rapamycin. To distinguish these possibilities, we first sub-selected VNC nuclei and examined *Fkbp12* expression across age and treatment conditions. Although average *Fkbp12* expression was low in the full VNC population, *Fkbp12* was enriched in a distinct VNC subpopulation that emerged during aging (**Figures 4a** and **Extended Data Fig. 6a**). This aged VNC population also showed elevated expression of glycolysis and lipid-synthesis genes, suggesting that *Fkbp12* enrichment coincides with mTORC1-linked metabolic transcriptional programs in a subset of VNC nuclei (**Figures 4b** and **Extended Data Fig. 6b**).

**Figure 4.**
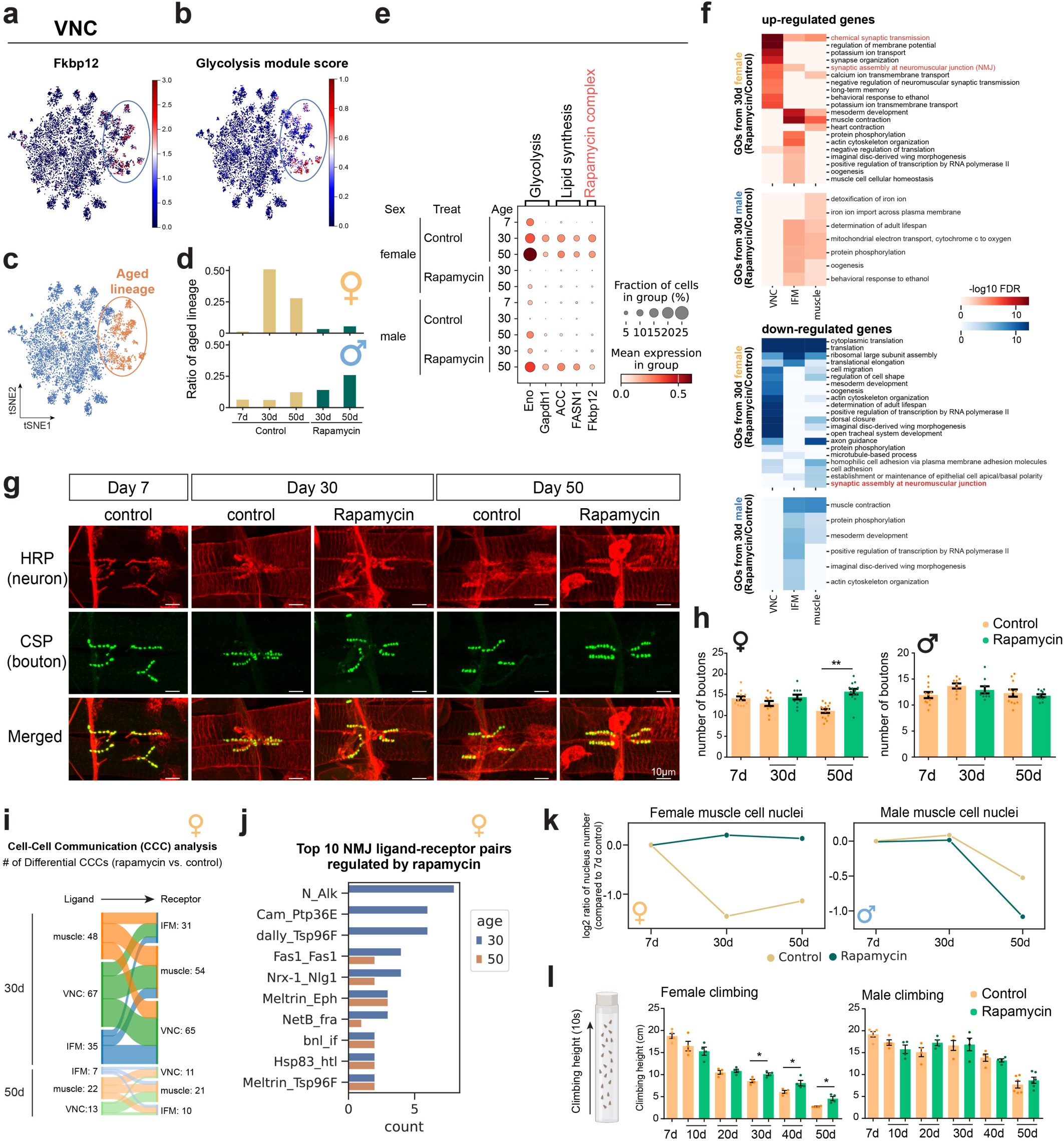
Rapamycin attenuates female-biased VNC and neuromuscular aging programs **a.** tSNE visualization of VNC nuclei showing *Fkbp12* expression. **b.** tSNE visualization of glycolysis module scores in VNC nuclei. **c.** Definition of the aged VNC lineage based on the glycolysis-high population. **d.** Quantification of aged-lineage ratios in female and male VNC nuclei across age and treatment conditions. **e.** Dot plots showing expression of glycolysis genes, lipid-synthesis genes, and *Fkbp12* in VNC nuclei across sex, age, and treatment conditions. **f.** GO analysis of rapamycin-induced DEGs in VNC, indirect flight muscle, and muscle cells from females and males. **g.** Representative confocal images of adult abdominal neuromuscular junctions from control and rapamycin-treated females across ages. **h.** Quantification of bouton number per muscle fiber in females and males across age and treatment conditions. Total quantified bouton numbers are 357∼596 from 28∼40 muscle fibers of each sample. Bars indicate mean ± SEM; **: unpaired t-test, p ≤ 0.01. **i.** Cell-cell communication analysis among VNC, indirect flight muscle, and muscle cells using FlyPhoneDB2. Differential ligand-receptor interactions between rapamycin-treated and control samples are shown for female flies at day 30 and day 50. **j.** Top 10 ligand-receptor pairs regulated by rapamycin between VNC, muscle, and IFM compartments in female flies at day 30 and day 50. **k.** Changes in muscle nucleus ratios across age and treatment conditions. **l.** Climbing performance across age and treatment conditions in females and males. Bars indicate mean ± SEM; *p ≤ 0.05, p values were from parametric unpaired t test.

We therefore defined this glycolysis-high VNC population as an aging-associated lineage and quantified its abundance across sexes, ages, and treatments (**Figure 4c**). In females, the aged VNC lineage increased with age and was reduced by rapamycin treatment. In contrast, the corresponding population in males did not show a comparable age-associated increase and was not reduced by rapamycin (**Figure 4d**). Consistently, *Fkbp12* and mTORC1-linked metabolic genes increased with age and decreased after rapamycin treatment in female VNC nuclei, whereas these changes were weaker or absent in males (**Figures 4e** and **Extended Data Fig. 6c**). These results suggest that rapamycin preferentially attenuates an aging-associated, *Fkbp12*-enriched metabolic state in the female VNC.

Muscle cells also showed high rapamycin-induced DEG numbers despite low average *Fkbp12* expression. Unlike the VNC, however, muscle nuclei showed limited age- or treatment-associated changes in glycolysis and lipid-synthesis gene expression (**Extended Data Fig. 6d**). This suggested that rapamycin-associated transcriptomic remodeling in muscle may not primarily reflect direct suppression of a muscle-intrinsic mTORC1-linked metabolic program. To identify alternative mechanisms, we performed Gene Ontology (GO) analysis of rapamycin-induced DEGs in muscle and VNC. In females, but not males, rapamycin-responsive genes were enriched for synaptic and neuromuscular junction (NMJ)-associated terms across VNC, IFM, and muscle cells (**Figure 4f**). These results suggested that rapamycin may influence muscle aging in females through maintenance of neuromuscular connectivity.

Consistent with this possibility, adult abdominal NMJ staining showed that rapamycin-treated females retained higher bouton numbers than age-matched controls, whereas males showed no significant difference (**Figures 4g–h**). We next used FlyPhoneDB2, a cell-cell communication analysis tool, to examine potential ligand-receptor interactions among VNC, IFM, and muscle cells (Qadiri et al., 2025). This analysis identified 150 differential communications in 30-day females and 42 differential communications in 50-day females after rapamycin treatment (**Figure 4i** and **Supplementary Table 1**). From these interactions, we identified the top 10 ligand-receptor pairs across neuromuscular cell types (**Figure 4j**). Notably, the well-characterized NMJ ligand-receptor pair Neurexin-1 (Nrx-1)-Neuroligin-1 (Nlg1) (Banovic et al., 2010) showed increased interaction scores and higher expression in rapamycin-treated 30-day females, but not in males (**Extended Data Fig. 6e**). These findings support a female-biased effect of rapamycin on molecular programs associated with NMJ maintenance.

We finally asked whether the rapamycin-associated preservation of neuromuscular features was accompanied by a broader suppression of muscle aging. Consistent with our previous AFCA analysis, muscle nuclei declined with age in control flies of both sexes; however, the kinetics of this cellular attrition were profoundly sexually dimorphic. Females exhibited a significant reduction in muscle nuclei after day 30 (**Figure 4k**). Rapamycin successfully prevented this decline, a cellular preservation that translated into significantly improved climbing ability in aged females (**Figure 4l**). In contrast, male flies maintained stable muscle nuclei counts and climbing performance through day 30 before experiencing a sharp decline at a later age. Consequently, the feature of male aging masked early-life functional benefits of rapamycin in males (**Figures 4k and 4l**). Collectively, these results reveal a fundamentally sex-specific mechanism of pharmacological geroprotection: rapamycin rescues female neuromuscular aging through the synergistic suppression of an *Fkbp12*-enriched, mTORC1-linked metabolic state in the VNC and the robust preservation of programs governing NMJ integrity and motor function.

### Rapamycin mitigates the Convergent Aging Trajectory (CAT)

The presence of rapamycin-sensitive aged populations in reproductive cells and the VNC raised the possibility that these populations may share a common aging-associated transcriptional program. To test this idea, we identified genes differentially expressed between aged and non-aged populations within each of the three previously defined aged lineages: female germline cells, male accessory gland main cells, and VNC nuclei (**Figure 5a**). Intersecting these lineage-specific signatures identified 469 genes that were consistently altered across all three aged populations, including 291 commonly upregulated genes and 178 commonly downregulated genes (**Figure 5b** and **Extended Data Fig. 7a**). GO analysis showed that commonly upregulated genes were enriched for glycolysis, cytoplasmic translation, mitochondrial electron transport, proteostasis, and superoxide radical removal, whereas commonly downregulated genes were enriched for protein phosphorylation, mRNA splicing, and cellular morphology-related processes (**Figure 5b**). These shared signatures suggest that aged nuclei from distinct tissues converge toward a common transcriptional state that extends beyond glycolysis alone.

**Figure 5.**
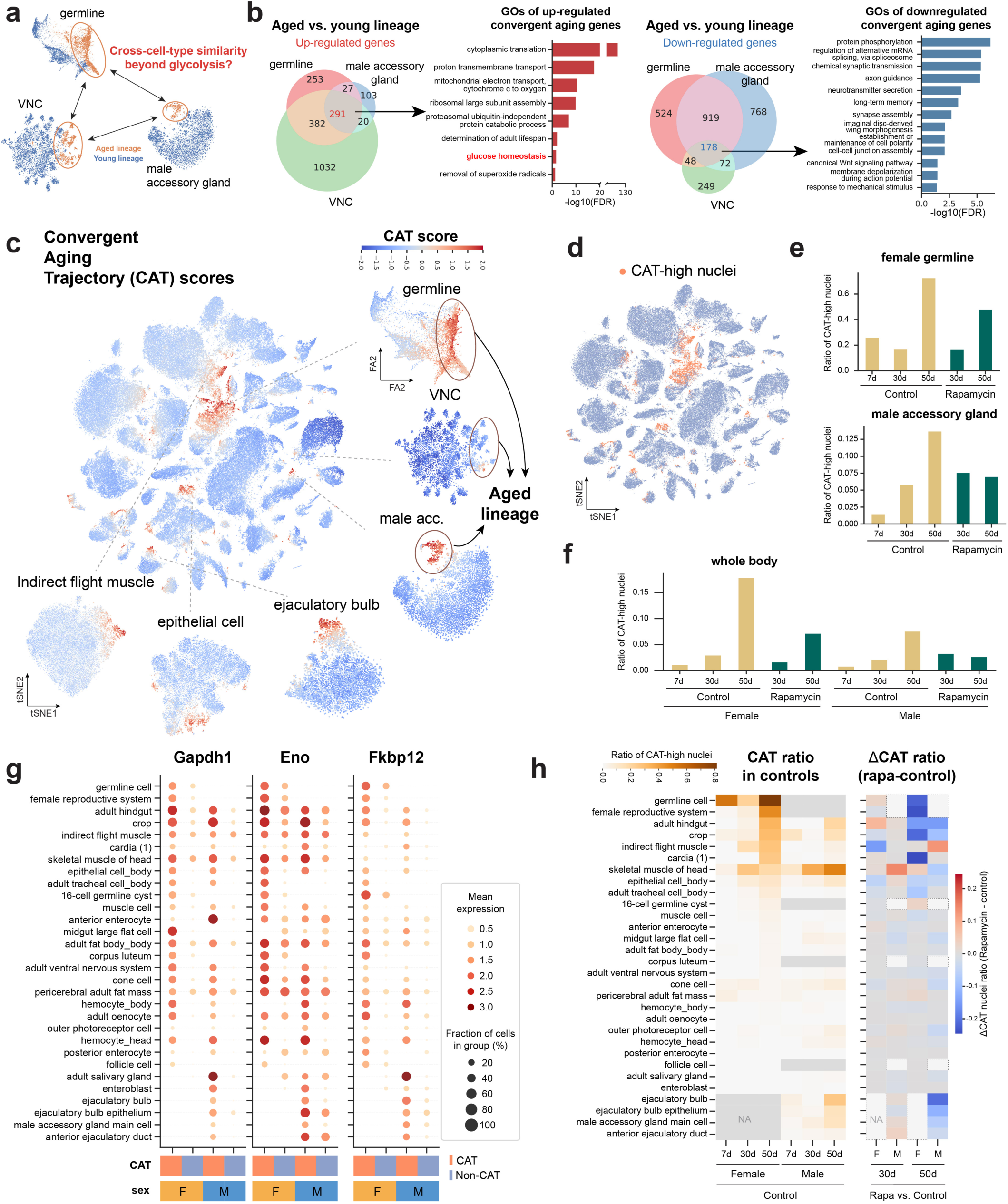
Accumulation of rapamycin-sensitive CAT-high nuclei during aging **a.** A highlight of aged lineages across three representative cell types: female germline, male accessory gland main cells, and VNC. **b.** Identification of a shared CAT gene signature. Venn diagrams summarize genes consistently upregulated or downregulated in aged lineages across the three representative cell types. GO enrichment analysis highlights biological processes enriched among commonly upregulated and downregulated genes. **c.** tSNE visualization of CAT scores across the body dataset with several cell types highlighted. **d.** tSNE visualization of CAT-high nuclei across the whole-body atlas. **e.** Ratios of CAT-high nuclei in female germline and male accessory gland main cells across age and treatment conditions. **f.** Whole-body CAT-high nucleus ratios across sex, age, and treatment conditions. **g.** Expression of glycolysis genes and *Fkbp12* in CAT-high and non-CAT nuclei across cell types and sexes. **h.** Heatmap summarizing CAT-high nucleus ratios in control samples and rapamycin-associated changes in CAT-high ratios across cell types, ages, and sexes.

We next asked whether this shared transcriptional program could identify similar aging-associated nuclei across the whole organism. Using the 469 shared DEGs, we defined a recurrent aging-associated transcriptional state as a Convergent Aging Trajectory (CAT). For each cell, we computed a CAT score based on the combined enrichment of commonly upregulated and downregulated gene sets (Methods and **Extended Data Fig. 7b–c**). High CAT scores were detected not only in the aged lineages used to define the signature but also in additional cell types, including the ejaculatory bulb, epithelial cells, and IFMs (**Figure 5c**). We then defined CAT-high nuclei using the CAT-score distribution and quantified their abundance across ages, sexes, and treatments (**Figure 5d and Extended Data Fig. 7d**). In female germline cells and male accessory gland main cells, CAT-high nuclei increased with age and were reduced by rapamycin treatment, closely matching the manually defined aged-lineage patterns (**Figures 5e, 3c,** and **3h**). At the whole-organism level, CAT-high nuclei also accumulated with age and were reduced after rapamycin administration, with higher CAT-high ratios and stronger rapamycin-associated reductions in females than in males (**Figure 5f**).

To determine whether CAT-high nuclei were linked to mTORC1-associated metabolic programs, we compared glycolysis gene expression between CAT-high and non-CAT nuclei across cell types. CAT-high nuclei showed higher glycolysis gene expression than non-CAT nuclei across diverse cell types, and they also exhibited elevated *Fkbp12* expression (**Figure 5g and Extended Data Fig. 7e–f**). These features are consistent with the earlier observation that rapamycin-responsive aged populations are enriched for mTORC1-linked transcriptional programs and suggest that CAT-high nuclei represent an aging-associated state with elevated rapamycin sensitivity. The prevalence of CAT-high nuclei varied substantially across cell types, with reproductive, digestive, and muscle-associated cell types generally showing higher CAT-high ratios (**Figure 5h**). However, rapamycin-mediated changes in CAT abundance also varied across tissues: reproductive and gut cell types showed marked reductions in CAT-high nuclei, whereas skeletal muscle showed more modest changes. Together, these results define CAT as a shared, rapamycin-sensitive aging-associated transcriptional state that emerges across multiple cell types and provides a quantitative framework for measuring mTORC1-linked aging and rapamycin-mediated geroprotection at single-nucleus resolution.

### The Convergent Aging Trajectory is reproducible in an independent aging atlas

To evaluate whether CAT-high nuclei represent a reproducible aging-associated state rather than a dataset-specific feature of Rapa-FCA, we applied the CAT signature to control samples from our previously generated Alzheimer’s disease Fly Cell Atlas (AD-FCA) dataset (Park et al., 2025). Although AD-FCA differs from Rapa-FCA in genetic background, culture temperature, and lifespan dynamics, CAT-high nuclei were again detected across multiple cell types and increased with age in control flies (**Extended Data Fig. 8**). This pattern was particularly evident in male reproductive cell types, including the accessory gland and ejaculatory bulb, consistent with the age-associated CAT accumulation observed in Rapa-FCA (**Figure 5h** and **Extended Data Fig. 8**). Sex differences were also observed in AD-FCA, with females showing higher CAT-high ratios and earlier age-associated increases than males. Together, these results indicate that CAT-high nuclei are reproducibly detected across independent whole-organism aging datasets, supporting CAT as a general aging-associated transcriptional state rather than a feature restricted to rapamycin-treated experiments.

### Multimodal quantification of sex- and cell-type-specific geroprotective effects

Although the CAT-high nucleus ratio provides a quantitative readout of rapamycin-sensitive, mTORC1-linked aging states, rapamycin may also mitigate cellular aging through mechanisms not fully captured by CAT abundance. We therefore integrated the CAT-based measurement established above with two additional readouts of cell-type-level aging: transcriptomic aging-clock predictions and recovered nucleus ratios (Buckley et al., 2023; Lu et al., 2023). For each measurement, aging-associated changes (aging effects) were first defined from control samples across age, and rapamycin-associated mitigation (geroprotection effects) were then estimated by comparing rapamycin-treated samples with age-matched controls (**Figure 6a**). This framework allowed us to compare whether rapamycin shifted each cell type away from, or toward its normal aging-associated trajectory.

**Figure 6.**
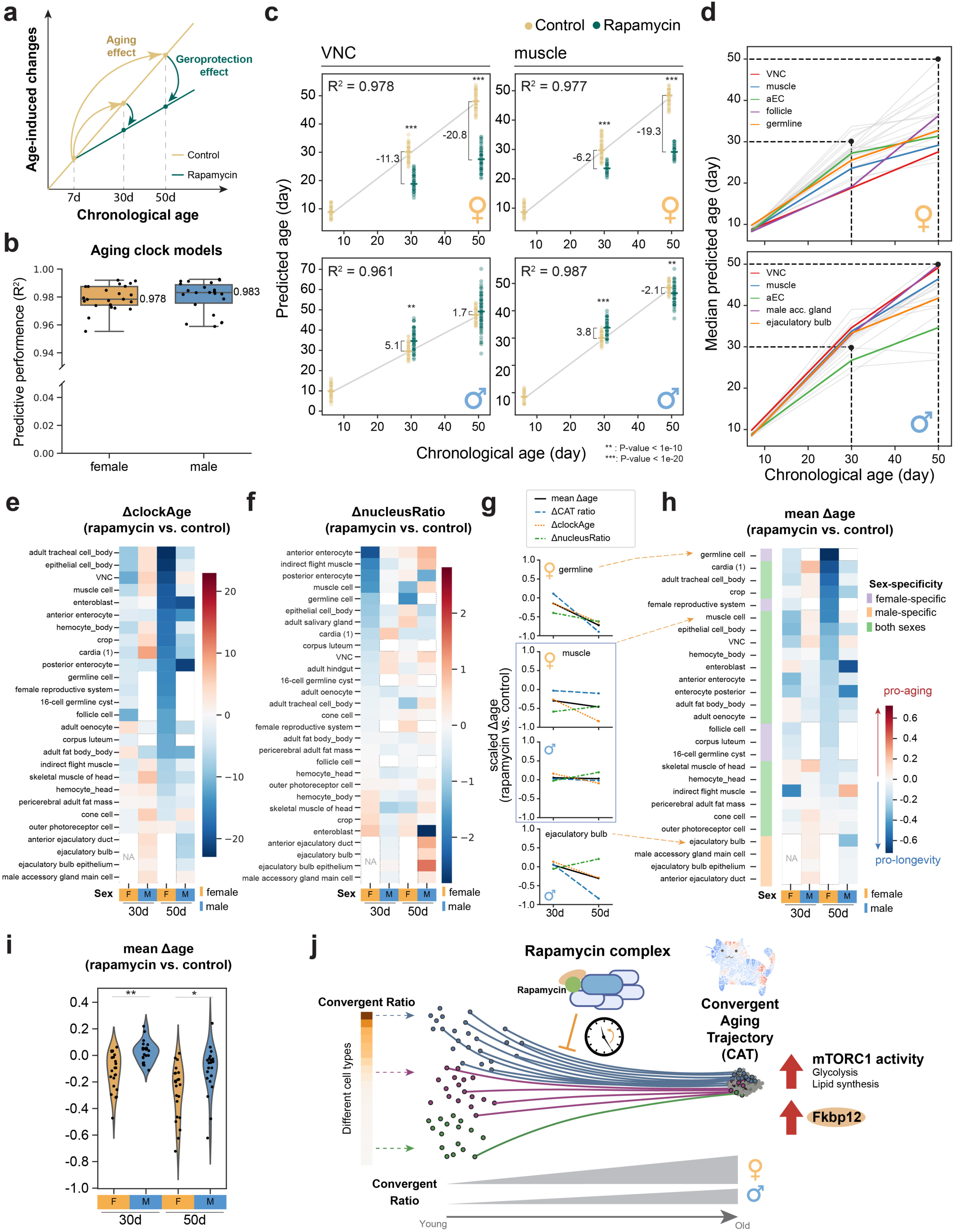
Systematic measurement of rapamycin-mediated geroprotection across the whole organism **a.** Schematic illustrating the calculation of aging effects and geroprotection effects. **b.** Predictive performance of cell-type-specific transcriptomic aging-clock models trained on pseudobulk profiles from control samples. **c.** Predicted transcriptomic ages of VNC and muscle cells from control and rapamycin-treated samples. The difference in predicted age between rapamycin-treated and control samples was assessed using the Wilcoxon rank-sum test. **d.** Median predicted transcriptomic ages for selected cell types across sex, age, and treatment conditions, highlighting cell-type- and sex-dependent responses to rapamycin. **e.** Heatmap of ΔclockAge across cell types, ages, and sexes. **f.** Heatmap of ΔnucleusRatio across cell types, ages, and sexes. **g.** Comparison of three scaled geroprotection measurements, ΔCAT ratio, ΔclockAge, and ΔnucleusRatio, in representative cell types. **h.** Heatmap of the integrated Δage score across cell types, ages, and sexes. Blue indicates stronger rapamycin-associated geroprotection, whereas red indicates limited or potentially adverse age-associated shifts. **i.** Violin plots showing the global distribution of integrated Δage scores across the cell-type atlas in females and males. Wilcoxon rank-sum test; P < 0.01 (*), P < 0.001 (**). **j.** Working model of sex-specific CAT accumulation and rapamycin-mediated CAT suppression. Female flies show broader accumulation of rapamycin-sensitive CAT-high nuclei and stronger CAT mitigation after rapamycin treatment, whereas male responses are more temporally and cell-type dependent.

To predict the biological age, we trained cell-type-specific aging-clock models using pseudobulk profiles from control samples (Buckley et al., 2023). These models showed strong predictive performance across cell types in both females and males (**Figure 6b**). We then applied the trained models to rapamycin-treated samples and calculated ΔclockAge as the difference in predicted age between rapamycin-treated and age-matched control samples, with negative values indicating a younger predicted age after rapamycin treatment. This analysis revealed substantial sex differences in rapamycin-associated age shifts. In female VNC and muscle cells, rapamycin reduced predicted age across both examined treatment ages, whereas the corresponding male cell types showed marginal or even age-increasing effects (**Figure 6c**). This female-biased reduction in predicted age was also observed across additional cell types when comparing median predicted ages and ΔclockAge values (**Figure 6d–e** and **Extended Data Fig. 9a**). Notably, many male cell types showed increased predicted age at day 30 after rapamycin treatment, whereas most cell types showed reduced predicted age at day 50, suggesting that rapamycin-associated geroprotection in males may be more age-dependent than in females.

We next used recovered nucleus ratios as an orthogonal readout of cell-type-level aging features. For each cell type, we calculated ΔnucleusRatio as the difference in recovered nucleus ratio between rapamycin-treated and age-matched control samples, and interpreted this change relative to the direction of age-associated shifts observed in control flies (**Figure 6f**, see **Methods**). Across cell types, ΔnucleusRatio suggested stronger rapamycin-associated effects at day 30 than at day 50, particularly in females (**Extended Data Fig. 9B**). The strongest effects in 30-day females were observed in enterocyte, muscle, and germline populations. Among these, germline and muscle cells showed consistent rapamycin-associated mitigation across both ages, supporting their sustained responsiveness to rapamycin.

Finally, we integrated the three geroprotection measurements to compare rapamycin effects across cell types, sexes, and ages. Each metric was scaled to a common range, allowing comparison of CAT reduction, transcriptomic age shifts, and nucleus-ratio changes within the same framework (**Figure 6f–i** and **Extended Data Fig. 9c**). This analysis revealed distinct modes of rapamycin-associated aging mitigation. Female germline cells showed strong geroprotective effects across all three measurements, indicating broad rapamycin sensitivity. Female muscle cells showed limited CAT reduction but clear mitigation based on transcriptomic age and nucleus-ratio measurements, consistent with the neuromuscular effects described above. In contrast, male ejaculatory bulb cells showed reduced CAT-high nuclei and transcriptomic age but little change in nucleus ratio, suggesting that different readouts capture different aspects of rapamycin response.

Overall, the integrated Δage score identified three major patterns: several cell types in both sexes, such as digestive cell types, and reproductive cell types showed consistent geroprotective responses; females exhibited stronger rapamycin-associated benefits than males across multiple measurements; and early rapamycin treatment produced limited or potentially unfavorable age-associated shifts in many male cell types, whereas female cell types showed more consistent protection across ages.

Taken together, the Rapa-FCA provides the first organism-wide, single-nucleus map of rapamycin action. By defining tissue-specific benefits, biological vulnerabilities, and sex-dependent responses, this study establishes a comprehensive framework to refine longevity interventions. These foundational insights and multimodal analytical strategies are broadly applicable to both basic and preclinical research and the wider single-cell transcriptomics community.

## Discussion

### CAT Defines a Rapamycin-Sensitive Convergent Aging State

Our Rapa-FCA reveals an unexpected layer of complexity in cellular aging. Although individual cell types undergo distinct aging trajectories, a subset of aged nuclei from diverse tissues converges toward a shared transcriptional state, which we term Convergent Aging Trajectory or CAT. CAT-high nuclei are enriched for mTORC1-linked transcriptional programs, including glycolysis and lipid metabolism, and are most abundant in tissues with high proliferative, reproductive, or metabolic demands, such as the reproductive and digestive systems. Importantly, the abundance of CAT-high nuclei is reduced in rapamycin-treated flies, suggesting that CAT represents a rapamycin-sensitive aging-associated state. This effect is more pronounced in females, consistent with the stronger transcriptomic and organismal responses to rapamycin observed in female flies.

Beyond canonical mTORC1-associated pathways, CAT-high nuclei exhibit additional shared transcriptional features across cell types (**Figure 5b-c**). These include changes in mitochondrial electron transport, cytoplasmic translation, proteostasis, and oxidative stress-response programs. These signatures suggest that CAT is not simply a glycolytic state, but rather a broader aging-associated cellular state involving metabolic rewiring and stress adaptation. Several lifespan-associated metabolites, including NAD+ and α-ketoglutarate, decline or become dysregulated during aging (Chin et al., 2014; Gomes et al., 2013; Massudi et al., 2012; Shahmirzadi et al., 2020), raising the possibility that CAT-high nuclei contribute to age-associated metabolic drift through coordinated changes in nutrient sensing, mitochondrial activity, and redox homeostasis. Future integration of metabolomic, proteomic, and functional assays will be important to determine whether CAT-high nuclei directly drive these metabolic changes or instead reflect cellular adaptation to age-associated stress.

The enrichment of *Fkbp12*, the primary binding partner required for rapamycin-mediated mTORC1 inhibition, provides a mechanistic link between CAT-high nuclei and rapamycin sensitivity. Across cell types, *Fkbp12* expression is positively associated with the magnitude of rapamycin-induced transcriptional responses, and CAT-high nuclei exhibit elevated *Fkbp12* together with mTORC1-linked metabolic programs. This coordinated enrichment suggests that CAT is not merely an aging-associated transcriptional state, but a cellular state poised for rapamycin responsiveness. In this model, aging promotes the emergence of metabolically active, *Fkbp12*-high nuclei across diverse tissues, thereby creating a shared molecular vulnerability that can be suppressed by rapamycin.

Together, these findings support a model in which rapamycin does not act uniformly across all cell types, but preferentially suppresses a convergent aging-associated cellular state marked by elevated *Fkbp12* expression and mTORC1-linked metabolic activity in the peripheral cell types. This model provides a potential explanation for how rapamycin can generate organism-wide geroprotective effects despite the diversity of tissues and aging trajectories: instead of targeting cell identity per se, rapamycin may act on a shared aging state that emerges repeatedly across susceptible cells. Thus, CAT provides both a mechanistic framework for understanding rapamycin-mediated longevity and a practical cellular-state readout for identifying rapamycin-sensitive aging populations across the organism.

### Determinants of Rapamycin Efficacy across Cell types

What factors can affect rapamycin’s efficacy on different cell types? First, tissue accessibility may influence the magnitude of rapamycin response. Rapamycin is a large (914.2 Da), hydrophobic compound with limited aqueous solubility, and mammalian studies suggest that rapamycin and its analogs can show limited blood-brain barrier penetration (Brandt et al., 2018; Gonzales et al., 2025; Song et al., 2025). Consistent with this possibility, Rapa-FCA revealed modest transcriptional changes in the brain nervous system, whereas peripheral cell types showed stronger remodeling. However, drug accessibility was not directly measured in this study. Future tissue-specific pharmacokinetic measurements will therefore be required to determine whether limited rapamycin exposure contributes to the reduced transcriptional responsiveness of specific tissues, including the head neuronal populations.

Second, rapamycin efficacy likely depends on the degree of Fkbp12-mTORC1 target engagement within each cellular state. Rapamycin suppresses mTORC1 through Fkbp12, and CAT-high nuclei show coordinated enrichment of *Fkbp12* and mTORC1-linked metabolic programs. Thus, cell types with higher CAT abundance, such as reproductive and digestive populations, may contain a larger fraction of cells with the molecular features associated with rapamycin responsiveness. In this context, CAT provides a measurable transcriptional readout for identifying cell types and cellular states most likely to respond to rapamycin.

Third, rapamycin efficacy may also depend on the relationship between CAT-high nuclei and other aging-associated features. CAT defines an intrinsic aging-associated state that is directly reduced by rapamycin in several cell types, but some protective effects are not fully explained by CAT reduction alone. Female muscle cells, for example, show limited CAT reduction but clear geroprotection based on transcriptomic age, nucleus-ratio preservation, NMJ-associated programs, and climbing performance. This pattern suggests that rapamycin may suppress CAT-high states in one cellular compartment, such as VNC, while indirectly benefiting connected tissues through altered neuromuscular or intercellular communication. Such CAT-associated crosstalk may be particularly important for aging features that emerge from tissue interactions rather than from a purely cell-autonomous state.

Together, these three layers, tissue accessibility, Fkbp12-mTORC1-associated cellular responsiveness, and CAT-associated crosstalk with other aging features, provide a framework for understanding why rapamycin produces strong geroprotection in some cell types but limited or uncoupled responses in others. This framework helps reconcile two observations from Rapa-FCA: rapamycin preferentially suppresses CAT-high states in several reproductive and digestive cell types, yet can also improve aging-associated features beyond CAT reduction.

### Systemic Sexual Dimorphism in Aging and Pro-longevity

Sex is a major determinant of aging trajectories and responses to geroprotective interventions (Jiang et al., 2025). Rapamycin provides a clear example: although it extends lifespan in both sexes, its benefits are often stronger in females. Previous studies have suggested several contributing mechanisms, including sex differences in drug exposure, baseline mTORC1 activity, intestinal homeostasis, autophagy, reproduction, and systemic metabolism (Bjedov et al., 2010; Fan et al., 2015; Harrison et al., 2009; Miller et al., 2014; Regan et al., 2016, 2022). Our Rapa-FCA extends these observations by showing that sexual dimorphism is embedded at the cellular-state level. Female flies accumulate higher levels of CAT-high nuclei across multiple tissues, and these nuclei are enriched for *Fkbp12* and mTORC1-linked metabolic programs. The preferential reduction of CAT-high nuclei in rapamycin-treated females suggests that female-biased rapamycin efficacy may arise, at least in part, from the greater accumulation of a rapamycin-sensitive aging state. Thus, females may benefit more strongly not simply because of global pharmacological differences, but because they harbor broader *Fkbp12*-high, mTORC1-linked cellular substrates for rapamycin action.

In contrast, male responses to rapamycin appear more temporally and cell-type dependent. Several male cell types show limited or even unfavorable transcriptomic responses at earlier ages, whereas stronger geroprotective effects emerge later in life. This pattern suggests that male aging may be less dominated by early CAT accumulation, may involve mechanisms less sensitive to mTORC1 inhibition, or may be influenced by male-specific reproductive and metabolic programs. More broadly, these findings may help explain why different longevity interventions show distinct sex biases: each intervention may act most effectively when its molecular target aligns with the aging-associated cellular states enriched in that sex. Rapa-FCA therefore provides a framework for moving beyond sex as a descriptive variable and toward identifying the sex-biased cellular states that shape intervention efficacy, an important step for precision geroscience and for age-related diseases with sex-biased incidence or progression.

### Precision Aging Intervention: From Molecular Pathways to Cellular States

Most geroprotective strategies have been developed around genes, pathways, or hallmarks that regulate aging, such as nutrient sensing, proteostasis, mitochondrial function, senescence, and stem-cell maintenance (Guarente et al., 2024; Kroemer et al., 2025). This pathway-centered framework has been highly productive, but aging is ultimately expressed through heterogeneous cellular changes across tissues. A single intervention may therefore produce beneficial, neutral, or adverse effects depending on which cell types or cellular states are present in a given organismal context. Our Rapa-FCA supports a complementary framework in which aging interventions are evaluated not only by their molecular targets, but also by the cellular states they modify. In this context, CAT provides a measurable aging-associated state that emerges across multiple cell types, is enriched for *Fkbp12* and mTORC1-linked metabolic programs, and is reduced by rapamycin treatment. Thus, CAT may serve as a cellular-state readout for identifying rapamycin-sensitive aging populations and for assessing where mTORC1 inhibition is most likely to produce geroprotective effects.

This cellular-state perspective may help refine precision geroscience. Rather than assuming that a pathway-targeting intervention acts uniformly across the organism, whole-organism single-cell profiling can reveal which aging-associated states are suppressed, unchanged, or potentially exacerbated by treatment. For rapamycin, the reduction of CAT-high nuclei suggests that targeting an mTORC1-linked cellular state can generate coordinated effects across diverse tissues, particularly in females. More broadly, similar analyses could be applied to other interventions to determine whether they act on CAT-like states, distinct sex-biased aging states, senescent-like populations, stem-cell states, or tissue-specific stress programs. Such comparisons may provide rational principles for combination therapies: interventions could be paired not simply because they target different pathways, but because they act on complementary aging-associated cellular states. In this way, Rapa-FCA establishes a framework for moving from pathway-based geroprotection toward cellular-state-guided intervention design.

### Rapa-FCA: A Resource for Studying Rapamycin-associated Regulations

The conserved role of the rapamycin-mTORC1 axis in lifespan regulation underscores the importance of understanding its effects across the entire organism at cellular resolution. Rapa-FCA provides such a resource by integrating age, sex, tissue, cell type, and treatment information across more than 500,000 nuclei and 181 annotated cell types. This atlas enables systematic identification of rapamycin-responsive genes, sex-biased intervention effects, age-dependent treatment responses, and cell-type-specific aging trajectories. Beyond individual gene expression changes, Rapa-FCA provides a platform for studying how pharmacological intervention reshapes organism-wide cellular architecture, including direct suppression of rapamycin-sensitive aged states, indirect remodeling through intercellular communication, and tissue-specific limitations in intervention response. By revealing both the reach and the boundaries of rapamycin action, Rapa-FCA offers a foundation for designing more targeted, sex-aware, and cell-state-informed geroprotective strategies.

## Acknowledgments

We thank the members of the Bellen and Li laboratories for their insightful advice and discussions. We also thank Jiaye Chen for early assistance and discussions regarding the data portal. We thank Weiwei Dang for commenting on the manuscript. We acknowledge Shelby Yu-Hsiu Lu for her contribution to the design of the CAT symbol.

## Funding

H.L. is a CPRIT Scholar in Cancer Research (RR200063) and is supported by the NIH (U01 AG086143, DP2AT013275), the Longevity Impetus Grant, the Ted Nash Long Life Foundation, the Welch Foundation, and the Hevolution/AFAR Foundation. C.-Y.L., and A.-L.H are supported by NSTC 112-2926-I-A49A-503-G, 113-2926-I-A49A-502-G and AS-ASSA-113-03. T.J. is supported by the NIH/NIDDK F31DK141194-01A1 fellowship. Work in the NP laboratory is supported by NIH U01 AG086143 and the Howard Hughes Medical Institute. This project was supported by the Cytometry and Cell Sorting Core at Baylor College of Medicine with funding from the CPRIT Core Facility Support Award (CPRIT-RP240432) and the NIH (CA125123 and ODO36336), with the expert assistance of Joel M. Sederstrom.

## Author contributions

Conceptualization: T.-C.L., C.-Y.L., and H.L.; snRNA-seq: C.-Y.L., and Y.Q.; Computational analysis: T.-C.L., Z.Y., B.S., T.J., and M.Q.; Investigation: T.-C.L., C.-Y.L., Y.-J.P., N.A., E.H., N.P., A.-L.H., and H.L. Data portal and Rapa-FCA website: T.-C.L., and H.L.; Writing-original draft: T.-C.L., C.-Y.L., and H.L.; Writing-review and editing: all authors.; Supervision: H.L.; Funding acquisition: H.L.

## Competing interests

The authors declare no competing interests.

## Data, code, and materials availability

Raw FASTQ files, expression matrix, and processed h5ad files, including cell type annotations, are available at NCBI/GEO (accession number GEO: GSE322571). The code is available on github: https://github.com/MeshifLu/RapaFCA. All annotated data are also available for visualization, download, and custom analysis at https://hongjielilab.org/rapa-FCA/. Further information and requests for resources, materials and reagents should be directed to Hongjie Li.

## Materials and Method

### Fly sample and lifespan analysis

F1 flies of two wild type fly strains (virgin female w1118 and male OreR) were collected at day3 after first fly enclosed and maintained on the standard cornmeal-molasses medium (Li food) at 25C with 12-hour light-dark cycle for another 3 days for mating. Then male and female flies were separated into different cages (110 flies per cage) for aging. For lifespan analysis, the food was changed and dead flies were scored every 2-3 days. On day 7, the flies were applied to control food (DMSO, Dimethylsulfoxide, 0.1%) or Rapamycin food. The stock of rapamycin (LC Laboratories) was dissolved in DMSO (Dimethylsulfoxide) as 20 mM and added to the food to final concentration of 20 μM. Heads and bodies from flies at different ages were collected into a 1.5ml RNAase-free tube, snapped-frozen in liquid nitrogen and then stored at −80°C.

### Single-nucleus RNA sequencing (snRNA-seq)

The nuclei extraction was performed according to the previously described protocol (McLaughlin et al., 2022). BD FACSAria III sorter was used to collect Hoechst-positive nuclei, including all polyploidy nuclei. For each 10x Genomics run, 150-200k nuclei were sorted and centrifuged for 10 min at 1000g at 4°C. The nuclei were resuspended in 80 μl of 1x PBS with 0.5% BSA and RNase inhibitors. After counting with hemocytometers, 45-55k nuclei were loaded to achieve a target of 20k nuclei per sample. Libraries were generated using the Chromium Next GEM Single Cell 3’ HT Reagent Kits v3.1 (Dual Index) following 10x Genomics instructions. The final libraries were sequenced on an Illumina NovaSeq X Plus using PE150 (Novogene Cooperation Inc.) to a depth of approximately 30,000-50,000 reads per nucleus. To annotate cell identities, we integrated Rapa-FCA with our previously generated FCA, AFCA, and AD-FCA datasets, and transferred cell-type labels from AD-FCA to the rapamycin dataset (Li et al., 2022; Lu et al., 2023; Park et al., 2025). These annotations were then manually validated using established cell-type-specific markers.

### snRNA-seq data processing and quality control

The snRNA-seq data were processed using the computational pipeline established in our previous AFCA and AD-FCA studies (Lu et al., 2023; Park et al., 2025). Raw FASTQ files were aligned to the *Drosophila melanogaster* reference genome, FlyBase release 6.31, using Cell Ranger v6.1.2 with a pre-mRNA GTF file generated for the Fly Cell Atlas (Li et al., 2022). Ambient RNA contamination was removed using CellBender v0.2.2 (Fleming et al., 2023), and potential doublets were identified and removed using scDblFinder v1.8.0 (Germain et al., 2022).

Nuclei with fewer than 400 UMIs or fewer than 200 detected genes were removed. Genes detected in fewer than three nuclei were excluded from downstream analyses. Nuclei with gene counts or UMI counts greater than 10 median absolute deviations from the median, or with mitochondrial transcript fractions greater than 5%, were also removed. Downstream analyses were performed using Scanpy v1.8.2 (Wolf et al., 2018).

### Identification of rapamycin-responsive genes

To identify rapamycin-responsive genes, rapamycin-treated samples were compared with age-, sex-, tissue-, and cell-type-matched controls. Differential expression analysis was performed using the Wilcoxon rank-sum test implemented in Scanpy. Genes with a false discovery rate (FDR) below 0.05 were defined as rapamycin-responsive differentially expressed genes.

### Pathway enrichment and GO analysis

Pathway enrichment analysis was performed using the KEGG with *Drosophila*-specific pathway annotations (Kanehisa and Goto, 2000). Differentially expressed genes from each cell type were analyzed using GSEApy (Fang et al., 2023). To evaluate mTORC1-linked transcriptional programs, we focused on pathways expected to be suppressed by rapamycin-mediated mTORC1 inhibition, including glycolysis/gluconeogenesis, fatty acid biosynthesis, and biosynthesis of unsaturated fatty acids. Pathways associated with rapamycin-induced activation, including autophagy and lysosome pathways, were analyzed separately.

Glycolysis module scores were calculated using the score_genes() function in Scanpy. The glycolysis gene set was derived from genes enriched in the female germline glycolysis pathway and included *Eno*, *Gapdh1*, *CG32444*, *Pgm1*, *Pgi*, *Aldh*, *Pgk*, *Hex-A*, *Tpi*, *Ald1*, *AcCoAS*, *PyK*, *Pglym78*, and *Gapdh2*.

GO enrichment analysis was performed using GOATOOLS v1.2.3 (Klopfenstein et al., 2018). DEG lists from specific age, sex, treatment, and cell-type comparisons were tested for enriched GO Biological Process terms. The gene association file was retrieved from FlyBase FB2019_06.

### Germline snRNA-seq trajectory analysis

Female germline cells and 16-cell germline cyst cells were subsetted for trajectory analysis. Because of the complexity of germline subtypes, Rapa-FCA was integrated with the FCA, AFCA, and AD-FCA datasets to improve clustering and annotation resolution. Germline subtypes were annotated using marker-gene information from the FCA ovary dataset. Germline trajectories were analyzed using partition-based graph abstraction, PAGA, implemented in Scanpy (Wolf et al., 2018).

### Female fecundity

Female flies aged in lifespan cages containing either control food or food supplemented with rapamycin until the desired age were separated into vials containing standard lab food (N>35 per sample). The female was kept for egg laying for 3 days and then removed. The number of progeny in each vial was counted 12-15 days later. Females that died during the egg-laying period were excluded from the analysis.

### Male fecundity

Prior to mating, male flies were aged in lifespan cages containing either control food or food supplemented with rapamycin until the desired age. Individual male flies (N > 45 per sample) were then mated with two young wild-type female virgin flies in vials containing standard lab food. After mating at 25°C for 48 hours, the female flies were isolated into individual vials (one female per vial) and incubated at 25°C for 2 days to lay eggs. The mated females were then removed, and the number of progenies was recorded after 12-15 days at room temperature. Male fecundity was calculated as the average number of progenies from the two females. Any flies that died during the mating period were censored from the analysis.

### Cell-cell communication analysis among neuromuscular cell types

To examine rapamycin-associated changes in potential cell-cell communication among neuromuscular cell types, VNC, muscle cells, and indirect flight muscle cells were subsetted from the body dataset. Differential ligand-receptor interactions between rapamycin-treated and control flies were inferred using FlyPhoneDB2 (Qadiri et al., 2025). Ligand-receptor pairs with P < 0.05 or an absolute differential interaction score greater than 1.5 were considered differentially regulated after rapamycin treatment.

### Climbing assay

The climbing kit (Park et al., 2025) was used to measure the climbing ability of flies. Flies aged 7, 10, 20, 30, 40, or 50 days in lifespan cages were grouped into vials a day before the assay. For the climbing assay, flies were gently transferred to long vials in the climbing kit (20-25 flies per vial, total 80-100 files per sample). The climbing kit was gently tapped to force all flies to the bottom and then videotaped for 10 seconds. Flies climbed 5 times per set. Between each climbing, flies rested for 1 min. The climbing height of each animal at 10 seconds was measured by ImageJ. 4-5 sets of samples for each timepoint are plotted with GraphPad Prism.

### Adult neuromuscular junction (NMJ) staining and imaging

Fly abdomen filets (dorsal) were dissected in 1X PBS using Vannas Spring Scissors (FST, 15000-00) and forceps. The filets were placed into 8 strip 0.2ml PCR tubes, fixed with 4% PFA for 20 min at RT, and washed with 1X PBS. After fixation, the fat body tissues were carefully removed to reveal the dorsal muscles. After removal of fat body, filets were stained with mouse anti-CSP (DSHB, cat# DCSP-1, 1:20) in blocking buffer (5% NGS, 0.3% Triton X-100 in 1X PBS) at 4℃ overnight. Filets were thoroughly washed with 0.3% PBST, and stained with Alexa Fluor® 488 AffiniPure® Donkey Anti-Mouse IgG (H+L) (Jackson immunoResearch, cat# 715-545-151, 1:250) and Alexa Fluor® 594 AffiniPure™ Goat Anti-Horseradish Peroxidase (HRP) (Jackson immunoResearch, cat# 123-585-021, 1:250) in blocking buffer for 2 hours at RT. The nuclei were stained with DAPI (1:1000) for 10 min at RT. The filets were mounted in SlowFade™ Gold Antifade Mountant (Invitrogen, #S36936) and images were obtained with Leica STELLARIS 5 confocal microscope as Z series with 0.8um interval. Z series images were merged by ImageJ (Image-Stacks-Z projection-Max Intensity). NMJ analysis was carried out on A4 right-side hemi-segments. Number of synaptic boutons was quantified using presynaptic vesicle signals labelled by anti-CSP. Anti-HRP was used to visualize axons, and DAPI was used to confirm muscle fibers. At least 300 boutons were counted per sample. The mean bouton number per muscle fiber was calculated by dividing the total bouton count by the number of muscle fibers.

### Calculation of CAT scores and classification of CAT-high nuclei

Cell types containing distinct aged lineages were used to identify genes enriched or depleted in aged lineage relative to non-aged lineage. Differential expression analysis was performed using the Wilcoxon rank-sum test. Genes with FDR < 0.01 were defined as aged-lineage-specific DEGs. Upregulated and downregulated aged-lineage DEGs identified from female germline cells, male accessory gland main cells, and VNC nuclei were intersected to identify shared CAT-associated genes.

For each nucleus, log-normalized expression values of commonly upregulated and commonly downregulated CAT genes were averaged separately. The CAT score was calculated as the difference between the average expression of commonly upregulated genes and the average expression of commonly downregulated genes. CAT-score distributions were examined separately for body and head datasets across control and rapamycin-treated samples, sexes, and ages. Nuclei with CAT scores greater than the mean plus two standard deviations of the corresponding distribution were classified as CAT-high nuclei.

### Utilization of aging clock model for calculating the ΔclockAge

Cell-type-specific aging-clock models were trained using control samples with a pseudobulk BootstrapCell strategy (Buckley et al., 2023). For each cell type, sex and age group, UMI reads from randomly selected 15 nuclei or 5% of nuclei were summed to generate one pseudobulk BootstrapCell, followed by log normalization using Scanpy. This procedure was repeated 100 times for each cell type, sex and age group.

BootstrapCells generated from control samples were used to train Lasso regression models using the sklearn package. The Lasso penalty was fixed at alpha = 1.0 with max_iter = 10000 and tol = 1e-4. Model performance was evaluated using the coefficient of determination, R². After training, BootstrapCell expression profiles from control and rapamycin-treated samples were fitted to the corresponding cell-type-specific aging-clock model to obtain predicted ages. ΔclockAge was calculated as the difference between the median predicted age of rapamycin-treated samples and that of age- and sex-matched control samples. Negative ΔclockAge values indicate a younger predicted transcriptomic age after rapamycin treatment. Differences in predicted ages between control and rapamycin-treated samples were assessed using the Wilcoxon rank-sum test.

### Geroprotective effects on nucleus ratios measured by ΔnucleusRatio

Cell-type-specific nucleus ratios were calculated for each sample by dividing the number of nuclei assigned to a given cell type by the total number of nuclei recovered from the corresponding sample. Samples were analyzed separately by age, sex, treatment, and tissue compartment. For each cell type, the age-associated change in nucleus ratio was first defined in control flies by comparing aged control samples with the corresponding day 7 pretreatment baseline (aging effect, **Figure 6a**). Rapamycin-treated samples were normalized to the same day 7 baseline, and the rapamycin-associated change was then calculated by comparing the normalized rapamycin-treated value with the normalized age-matched control value (geroprotective effect, **Figure 6a**).

To determine whether this change represented a geroprotective or detrimental effect, the direction of the rapamycin-associated change was interpreted relative to the direction of aging observed in control flies. If rapamycin shifted the nucleus ratio in the opposite direction from the age-associated change in controls, the effect was considered geroprotective. If rapamycin shifted the nucleus ratio further in the same direction as the age-associated change in controls, the effect was considered age-enhancing or potentially detrimental. For example, because muscle nucleus ratios decline with age in control flies, a rapamycin-associated increase relative to age-matched controls was interpreted as protective in females (**Figure 4k**).

### Multimodal comparison of rapamycin’s geroprotective effects

To compare rapamycin-associated geroprotective effects measured by different approaches, ΔCAT ratio, ΔclockAge, and ΔnucleusRatio were scaled to a common range. For each measurement, the maximum absolute value was determined independently, and values were normalized using the maximum absolute value, resulting in scores ranging from −1 to 1. This scaling enabled cross-method comparison of rapamycin-associated changes across cell types, ages, and sexes. Integrated Δage scores were calculated by averaging the scaled values from the three measurements, with negative values indicating stronger rapamycin-associated geroprotection and positive values indicating limited or potentially adverse age-associated shifts.

## Notes

### Competing Interest Statement

The authors have declared no competing interest.

