## Supplemental Figures 1-9 for "Rapamycin Mitigates a Sex-biased Convergent Aging Trajectory"

#### Supplemental information

**Extended Data Figures 1–9, Supplementary Table 1, and supplemental references**

**Extended Data Fig. 1. Data quality and accessibility of Rapa-FCA**

- **a.** Overview of Rapa-FCA data accessibility. Processed data, visualization portals, downloadable files, raw sequencing data, and analysis code are available through the Li lab website, CELLxGENE, NCBI, and GitHub.
- **b.** tSNE visualizations of head and body datasets separated by age and treatment, showing the global distribution of nuclei across experimental conditions.
- **c.** Quality-control metrics for each Rapa-FCA sample, including the number of detected genes, UMI counts, and mitochondrial transcript ratios. Each point represents a nucleus, grouped by sample or library.

**Extended Data Fig. 2. Detailed cell-type annotations in the head dataset**

tSNE visualization of the Rapa-FCA head dataset colored by 107 detailed cell-type annotations.

**Extended Data Fig. 3. Detailed cell-type annotations in the body dataset**

tSNE visualization of the Rapa-FCA body dataset colored by 74 detailed cell-type annotations.

**Extended Data Fig. 4. *Fkbp12* expression is specifically associated with rapamycin-induced transcriptomic responses**

Comparison of the association between basal expression levels of FKBP family genes and rapamycin-induced DEG numbers across cell types. Significance values are shown as −log10(P values). *Fkbp12* shows a stronger association with the DEG number than most FKBP family members.

**Extended Data Fig. 5. Rapamycin attenuates mTORC1-linked aging programs in reproductive cells**

- **a.** Cell-type annotations and RNA velocity analysis of female germline nuclei across control and rapamycin-treated samples.
- **b.** Expression patterns of lipid-synthesis genes, including *Acc* and *FASN1*, in female germline nuclei.
- **c.** Age- and treatment-specific expression patterns of glycolysis genes, including *Eno*, *Gapdh1*, and *Tpi*, and lipid-synthesis genes, including *Acc* and *FASN1*, in male accessory gland main cells.
- **d.** tSNE visualization of glycolysis genes and lipid-synthesis genes in male accessory gland main cells.

**Extended Data Fig. 6. Aging- and rapamycin-associated changes in VNC, muscle, oenocyte, and fat body**

**a.** tSNE visualization of female VNC nuclei colored by age.
**b.** tSNE visualization of glycolysis genes and lipid-synthesis genes in VNC.
**c.** Expression and statistical comparisons of mTORC1-linked genes in female and male VNC nuclei across age and treatment conditions. Significance was assessed using the Wilcoxon rank-sum test and is shown as −log10(P value).
**d.** Expression patterns of glycolysis genes, lipid-synthesis genes, and *Fkbp12* in muscle cells across sex, age, and treatment conditions.
**e.** Rapamycin-associated changes in the Nrx-1-Nlg1 ligand-receptor pair between VNC and muscle/IFM compartments in female flies at day 30 and day 50. p values < 1e-5 from Wilcoxon Rank-Sum test.

**Extended Data Fig. 7. Definition and characterization of CAT-high nuclei**

**a.** Numbers of differentially expressed genes between aged and non-aged lineages in female germline cells, male accessory gland main cells, and VNC nuclei.
**b.** tSNE visualization showing the average expression of genes commonly upregulated across aged lineages.
**c.** tSNE visualization showing the average expression of genes commonly downregulated across aged lineages.
**d.** Distribution of CAT scores in body and head datasets. CAT-high nuclei were defined using a score threshold based on the CAT-score distribution, shown separately for body and head datasets.
**e.** Mean expression of representative CAT-associated genes, including *Gapdh1*, *Eno*, and *Fkbp12*, across cell types separated into CAT-high and non-CAT nuclei in both sexes.
**f.** Expression patterns of representative glycolysis genes and lipid-synthesis genes in CAT-high and non-CAT nuclei across cell types and sexes.

**Extended Data Fig. 8. CAT-high nuclei are reproducible in the independent AD-FCA dataset**

**a.** tSNE visualizations showing CAT-score distributions in selected AD-FCA control cell types, including male accessory gland, ejaculatory bulb, female germline, and whole-body datasets.
**b.** Quantification of CAT-high nucleus ratios in selected tissues and age groups from the AD-FCA body dataset.

**Extended Data Fig. 9. Quantification of rapamycin-associated geroprotection across cell types**

**a.** Violin plots showing the distribution of ΔclockAge across cell types and sexes.
**b.** Violin plots showing the distribution of ΔnucleusRatio across cell types and sexes.
**c.** Scaled geroprotection scores from three measurements, ΔCAT ratio, ΔclockAge, and ΔnucleusRatio, shown for representative cell types across age and sex. Scaling enables comparison of metric-specific similarities and differences in rapamycin-associated aging mitigation.

**Supplementary Table 1. Neuromuscular ligand-receptor pairs differentially regulated by rapamycin treatments in females.**

#### Extended Data Fig. 1

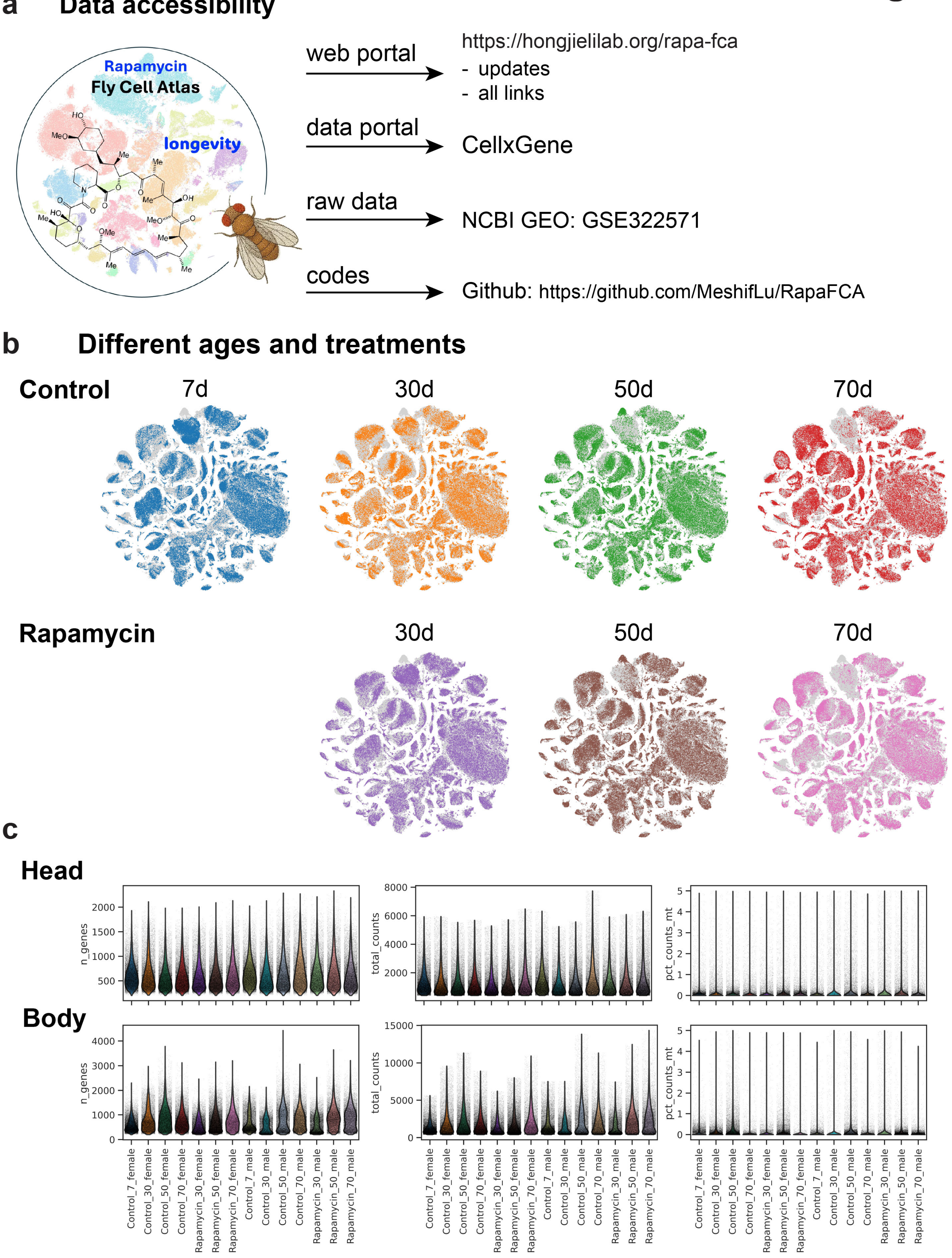

### Extended Data Fig. 2

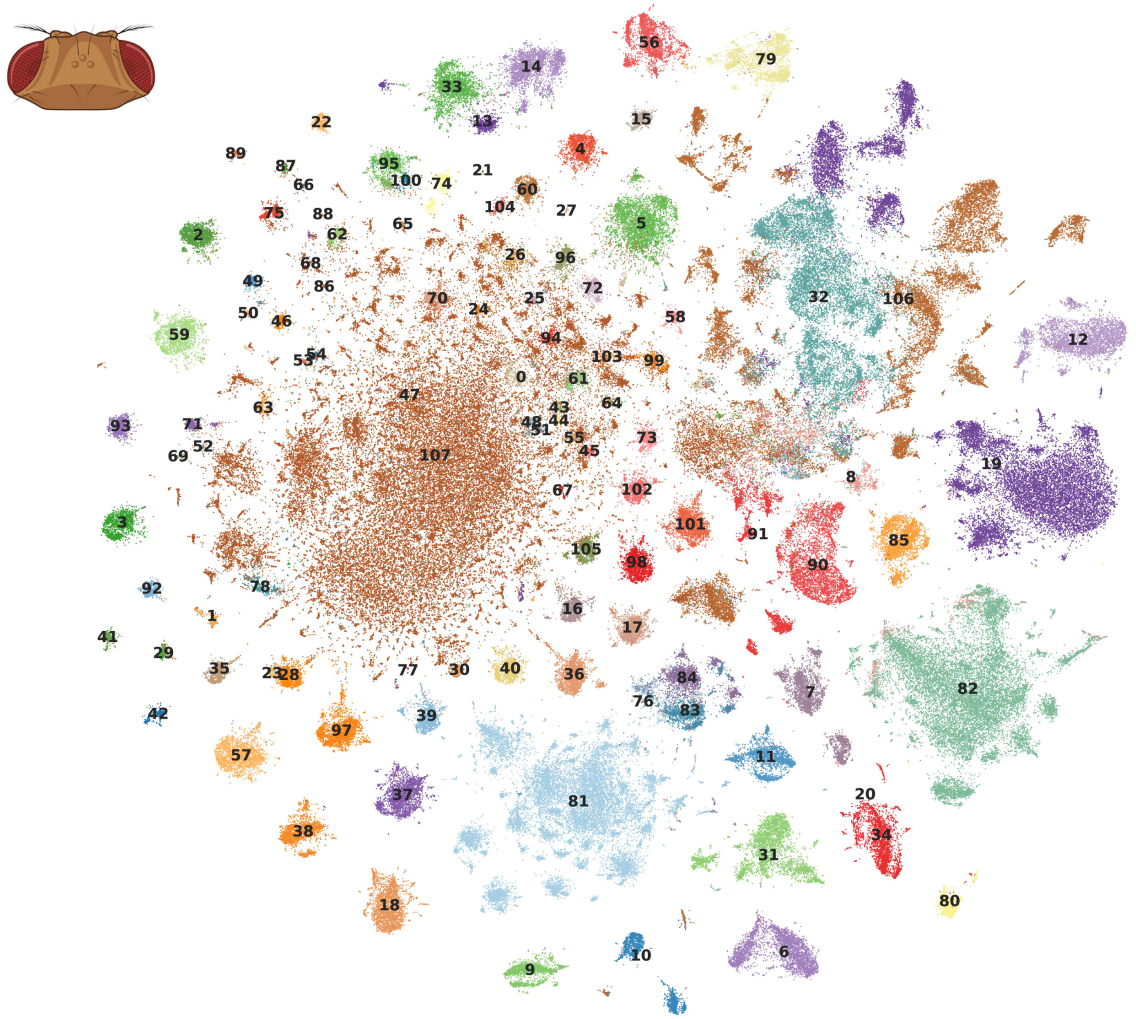

- 0.LDH-high cluster
- 1.Poxn neuron
- 2.T neuron T2
- 3.T neuron T2a
- 4.T neuron T3
- 5.T neuron T4/T5
- 6.adult brain cell body glial cell
- 7.adult brain perineurial glial cell
- 8.adult fat body_head
- 9.adult lamina epithelial/marginal glial cell
- 10.adult optic chiasma glial cell
- 11.adult reticular neuropil associated glial cell_head
- 12.adult tracheal cell_head
- 13.alpha'/beta' Kenyon cell
- 14.alpha/beta Kenyon cell
- 15.auditory sensory neuron
- 16.centrifugal neuron C2
- 17.centrifugal neuron C3
- 18.columnar neuron T1
- 19.cone cell
- 20.crystal cell_head
- 21.distal medullary amacrine neuron Dm1
- 22.distal medullary amacrine neuron Dm10
- 23.distal medullary amacrine neuron Dm11
- 24.distal medullary amacrine neuron Dm12
- 25.distal medullary amacrine neuron Dm2
- 26.distal medullary amacrine neuron Dm3
- 27.distal medullary amacrine neuron Dm4
- 28.distal medullary amacrine neuron Dm8
- 29.distal medullary amacrine neuron Dm9
- 30.dopaminergic PAM neuron
- 31.ensheathing glial cell
- 32.epithelial cell_head
- 33.gamma Kenyon cell
- 34.hemocyte_head
- 35.lamina intrinsic neuron
- 36.lamina monopolar neuron L1
- 37.lamina monopolar neuron L2
- 38.lamina monopolar neuron L3
- 39.lamina monopolar neuron L4
- 40.lamina monopolar neuron L5
- 41.lamina wide-field 1 neuron
- 42.lamina wide-field 2 neuron
- 43.lobula columnar neuron LC10a
- 44.lobula columnar neuron LC10b
- 45.lobula columnar neuron LC10d
- 46.lobula columnar neuron LC12
- 47.lobula columnar neuron LC14
- 48.lobula columnar neuron LC16
- 49.lobula columnar neuron LC17
- 50.lobula columnar neuron LC4
- 51.lobula columnar neuron LC6
- 52.lobula plate columnar LPC1
- 53.lobula plate-lobula columnar LPLC1
- 54.lobula plate-lobula columnar LPLC2
- 55.lobula-lobula plate columnar LLPC1
- 56.mechanosensory neuron
- 57.medullary intrinsic neuron Mi1
- 58.medullary intrinsic neuron Mi15
- 59.medullary intrinsic neuron Mi4
- 60.medullary intrinsic neuron Mi9
- 61.neuron 1
- 62.neuron 10
- 63.neuron 11
- 64.neuron 12
- 65.neuron 13
- 66.neuron 14
- 67.neuron 15
- 68.neuron 16
- 69.neuron 17
- 70.neuron 2
- 71.neuron 4
- 72.neuron 5
- 73.neuron 6
- 74.neuron 8
- 75.neuron 9
- 76.ocellus retinula cell
- 77.octopaminergic/tyraminergic neuron
- 78.olfactory projection neuron
- 79.olfactory receptor neuron
- 80.optic-lobe-associated cortex glial cell
- 81.outer photoreceptor cell
- 82.pericerebral adult fat mass
- 83.photoreceptor cell R7
- 84.photoreceptor cell R8
- 85.pigment cell
- 86.proximal medullary amacrine neuron Pm1
- 87.proximal medullary amacrine neuron Pm2
- 88.proximal medullary amacrine neuron Pm3*
- 89.proximal medullary amacrine neuron Pm4
- 90.skeletal muscle of head
- 91.subperineurial glial cell_head
- 92.transmedullary Y neuron TmY14
- 93.transmedullary Y neuron TmY3
- 94.transmedullary Y neuron TmY4
- 95.transmedullary Y neuron TmY5a
- 96.transmedullary Y neuron TmY8
- 97.transmedullary neuron Tm1
- 98.transmedullary neuron Tm2
- 99.transmedullary neuron Tm20
- 100.transmedullary neuron Tm29
- 101.transmedullary neuron Tm3a
- 102.transmedullary neuron Tm4
- 103.transmedullary neuron Tm5ab
- 104.transmedullary neuron Tm5c
- 105.transmedullary neuron Tm9
- 106.unannotated
- 107.uncharacterized CNS neuron

### Extended Data Fig. 3

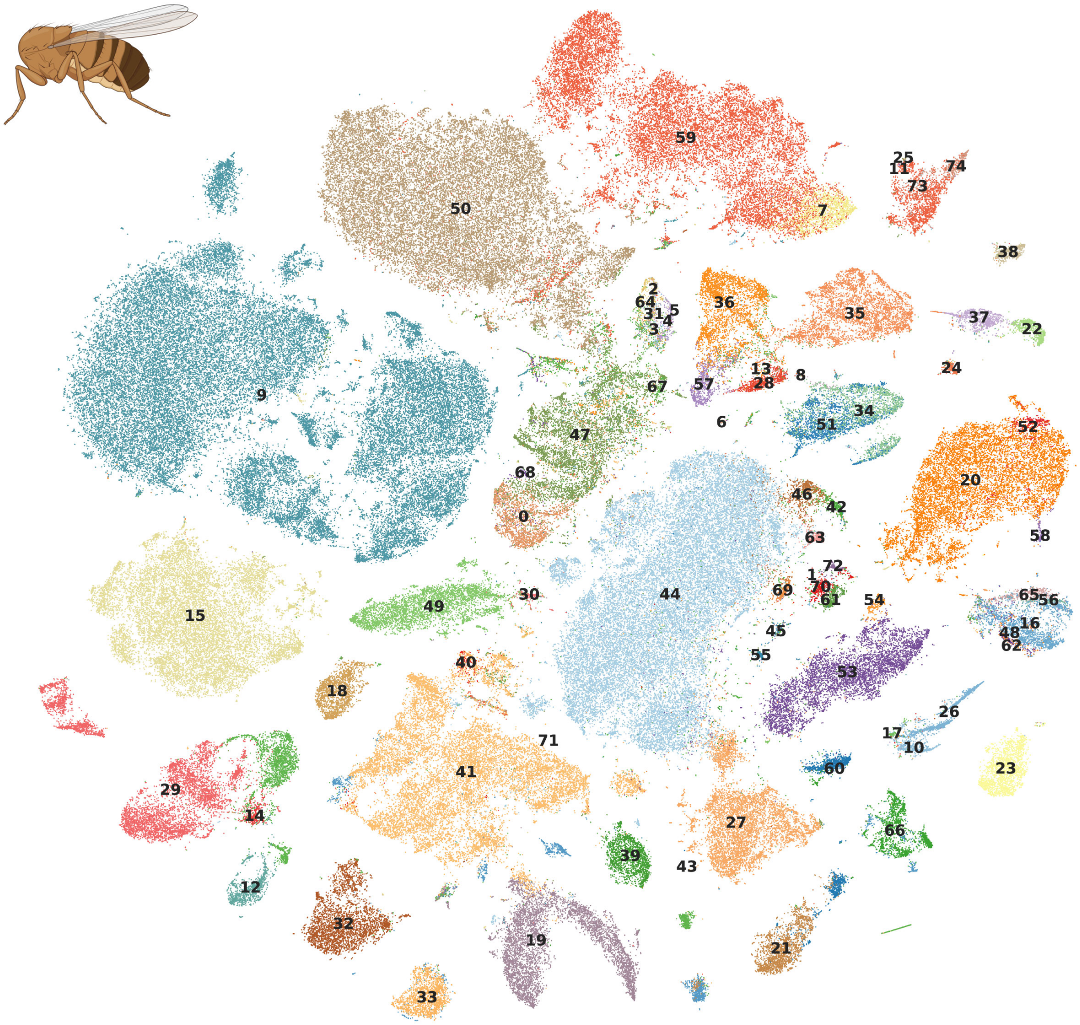

- 0.16-cell germline cyst in germarium region 2a and 2b
- 1.CNS surface associated glial cell
- 2.adult Malpighian tubule principal cell
- 3.adult Malpighian tubule principal cell of initial segment
- 4.adult Malpighian tubule principal cell of lower segment
- 5.adult Malpighian tubule principal cell of lower ureter
- 6.adult Malpighian tubule stellate cell of main segment
- 7.adult alary muscle
- 8.adult differentiating enterocyte
- 9.adult fat body_body
- 10.adult glial cell
- 11.adult heart ventral longitudinal muscle
- 12.adult hindgut
- 13.adult midgut enterocyte
- 14.adult midgut-hindgut hybrid zone
- 15.adult oenocyte
- 16.adult peripheral nervous system
- 17.adult reticular neuropil associated glial cell_body
- 18.adult salivary gland
- 19.adult tracheal cell_body
- 20.adult ventral nervous system
- 21.anterior ejaculatory duct
- 22.antimicrobial peptide-producing cell
- 23.cardia (1)
- 24.cardia (2)
- 25.cardiomyocyte, working adult heart muscle (non-ostia)
- 26.cell body glial cell
- 27.choriogenic main body follicle cell and corpus luteum
- 28.copper cell
- 29.crop
- 30.crystal cell_body
- 31.cyst cell
- 32.ejaculatory bulb
- 33.ejaculatory bulb epithelium
- 34.enteroblast
- 35.enterocyte of anterior adult midgut epithelium
- 36.enterocyte of posterior adult midgut epithelium
- 37.enterocyte-like
- 38.enteroendocrine cell
- 39.eo support cell
- 40.epidermal cell that specialized in antimicrobial response
- 41.epithelial cell_body
- 42.escort cell
- 43.female reproductive system
- 44.follicle cell
- 45.follicle cell uncharacterized
- 46.follicle stem cell and prefollicle cell
- 47.germline cell
- 48.gustatory receptor neuron
- 49.hemocyte_body
- 50.indirect flight muscle
- 51.intestinal stem cell
- 52.leg muscle motor neuron
- 53.male accessory gland main cell_roX1+
- 54.male accessory gland main cell_roX1-
- 55.male accessory gland secondary cell
- 56.mechanosensory neuron of haltere
- 57.midgut large flat cell
- 58.multidendritic neuron
- 59.muscle cell
- 60.oviduct
- 61.perineurial glial sheath
- 62.pheromone-sensing neuron
- 63.polar follicle cell
- 64.principal cell*
- 65.scolopidial neuron
- 66.seminal vesicle & testis epithelia
- 67.spermatid
- 68.spermatocyte
- 69.stalk follicle cell
- 70.subperineurial glial cell_body
- 71.unannotated
- 72.uncharacterized glial cell
- 73.visceral muscle of the crop
- 74.visceral muscle of the midgut

### Extended Data Fig. 4

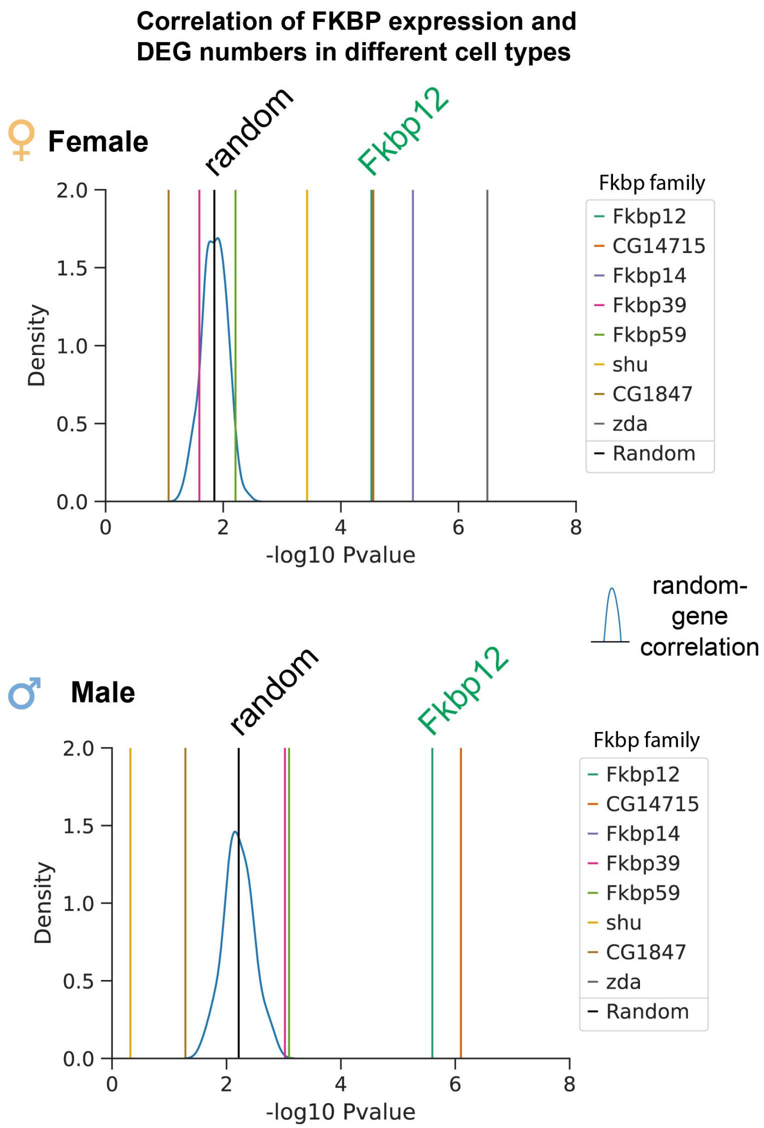

### Extended Data Fig. 5

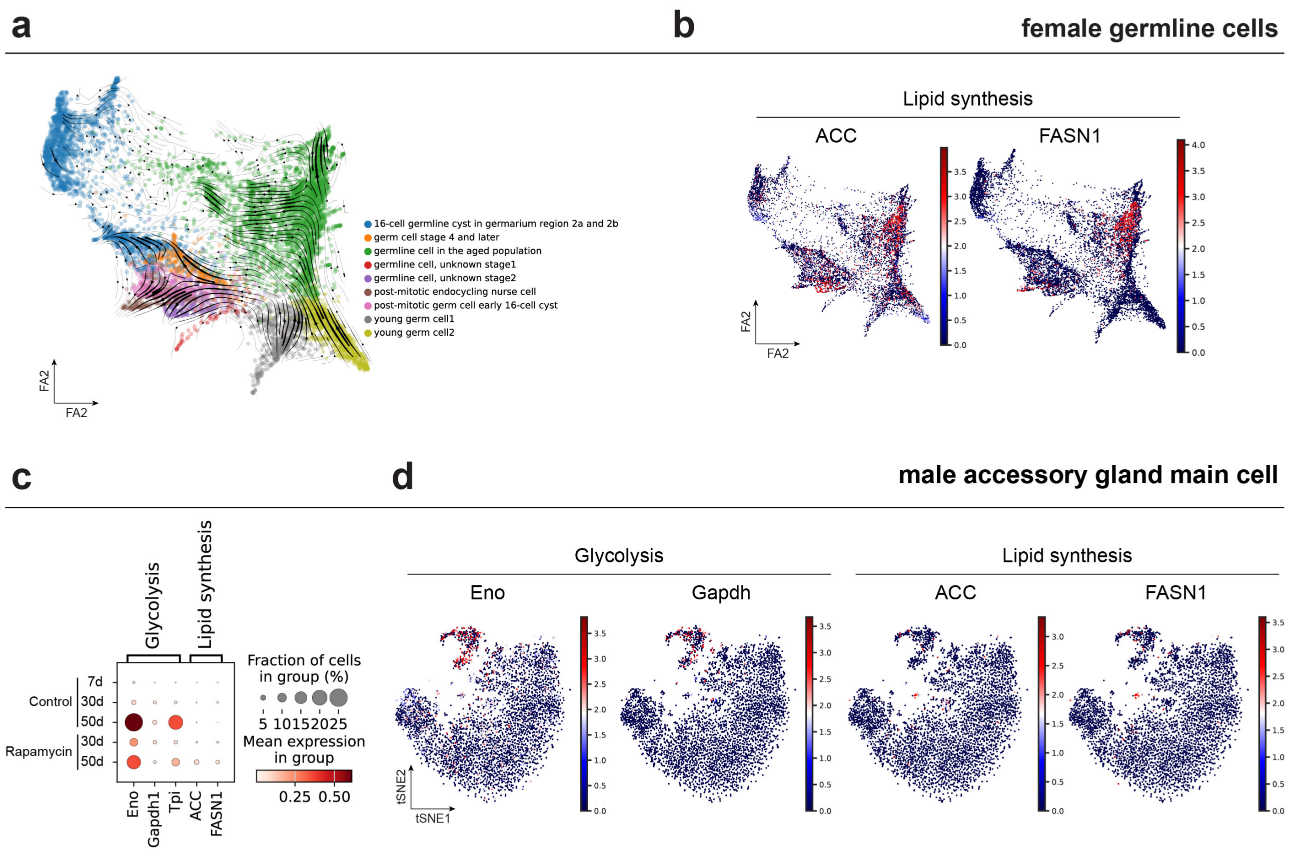

### Extended Data Fig. 6

**a**

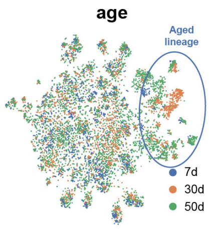


**b**

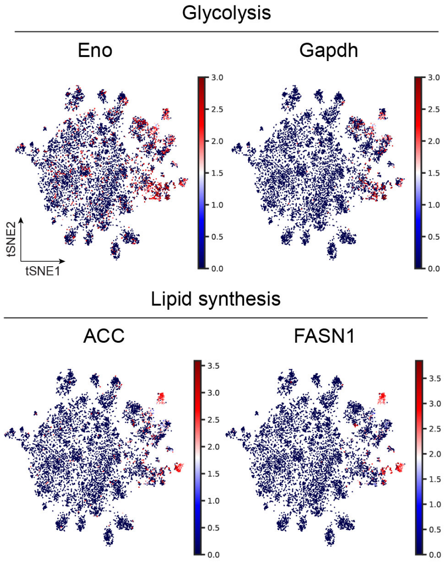


**c** **VNC**

Glycolysis

| Sex | Gene | Age | Padj | log2FC |
|---|---|---|---|---|
| female | Eno | 30 | 4.56E-05 | -5.260 |
| | | 50 | 3.60E-11 | -2.504 |
| | Gapdh1 | 30 | 0.03 | -5.500 |
| | | 50 | 1 | -2.399 |
| male | Eno | 30 | 1 | 2.847 |
| | | 50 | 0.151 | 0.682 |
| | Gapdh1 | 30 | 1 | 2.425 |
| | | 50 | 0.294 | 1.342 |

Rapamycin complex

| Sex | Gene | Age | Padj | log2FC |
|---|---|---|---|---|
| female | Fkbp12 | 30 | 0.021 | -3.734 |
| | | 50 | 0.105 | -2.772 |
| male | Fkbp12 | 30 | 1 | 25.922 |
| | | 50 | 0.767 | 1.612 |

Lipid synthesis

| Sex | Gene | Age | Padj | log2FC |
|---|---|---|---|---|
| female | ACC | 30 | 0.026 | -2.816 |
| | | 50 | 0.076 | -2.591 |
| | FASN1 | 30 | 0.199 | -3.530 |
| | | 50 | 0.258 | -3.101 |
| male | ACC | 30 | 1 | -0.602 |
| | | 50 | 0.009 | 1.988 |
| | FASN1 | 30 | 1 | 0.132 |
| | | 50 | 0.018 | 3.049 |

Wilcoxon Rank-Sum test
(Rapamycin vs. control)

**d** **Muscle cell**

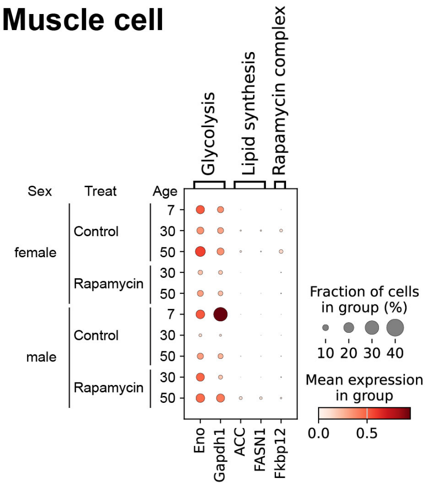


**e** **Differentially expressed ligand-receptor pair**

**NMJ: neuromuscular junction**

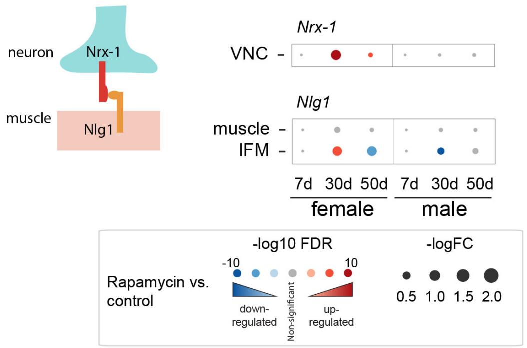

### Extended Data Fig. 7

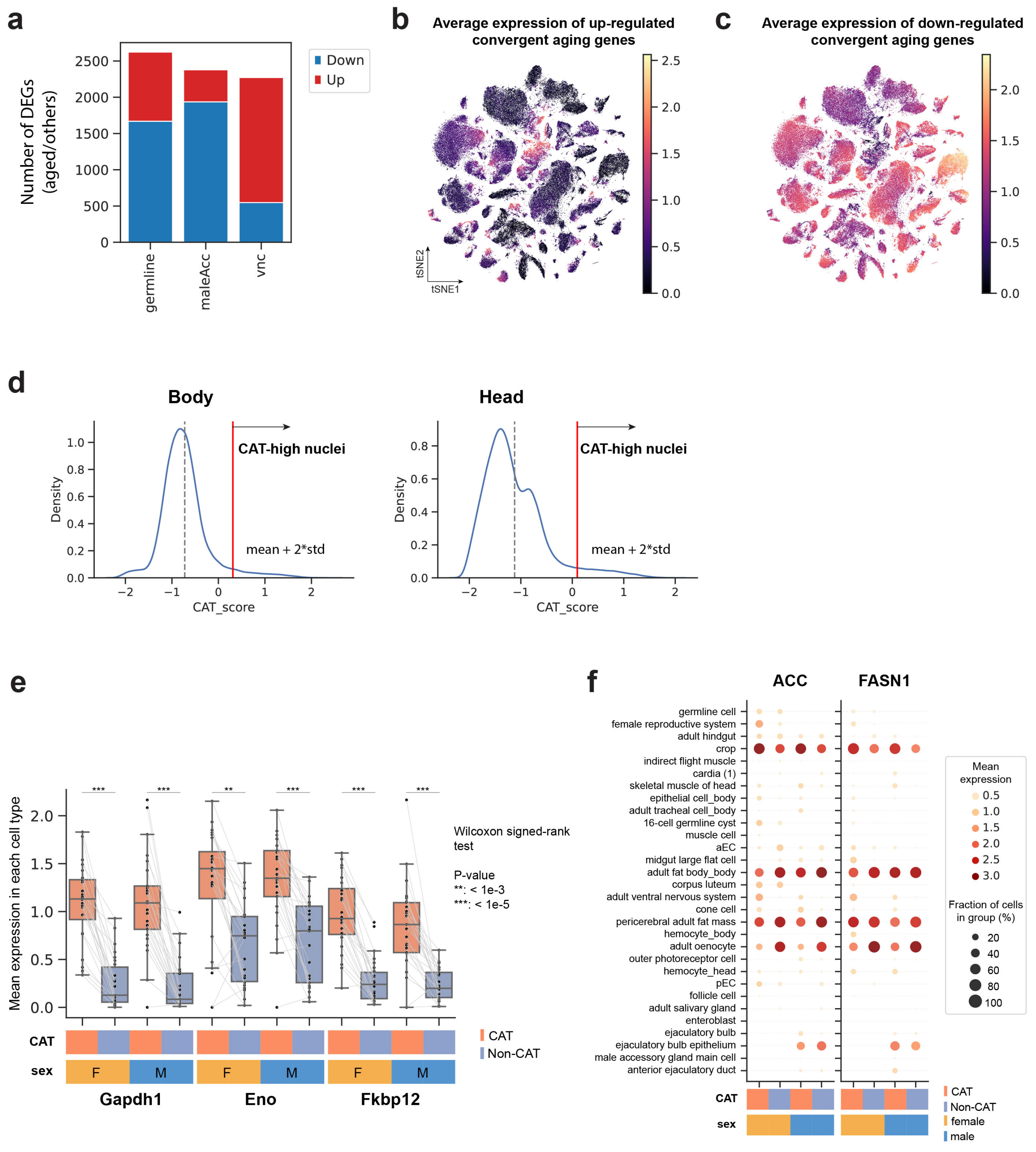

### Extended Data Fig. 8

**a** CAT scores from AD-FCA control body samples (29°C)

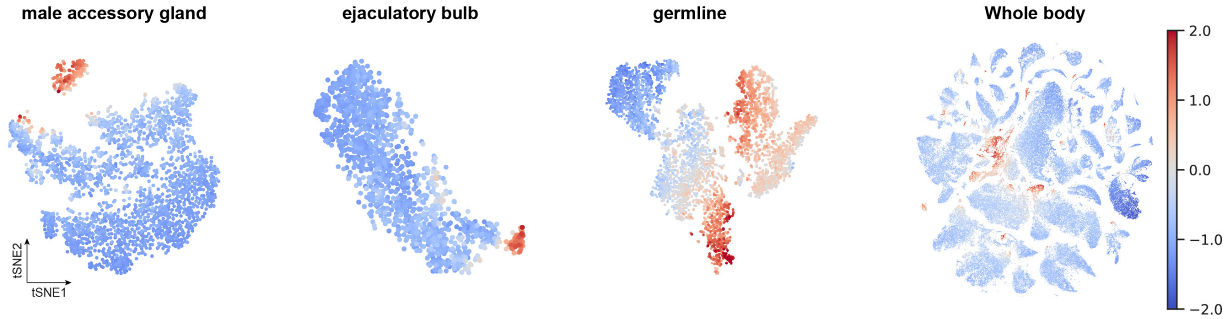


**b** Ratio of CAT-high nuclei from AD-FCA control body samples (29°C)

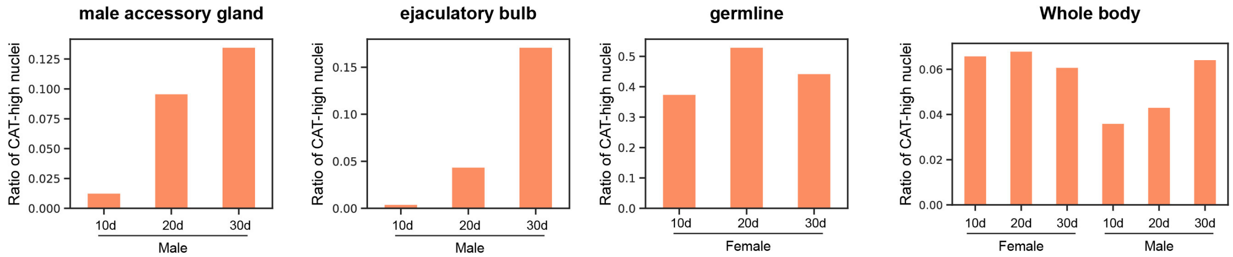

### Extended Data Fig. 9

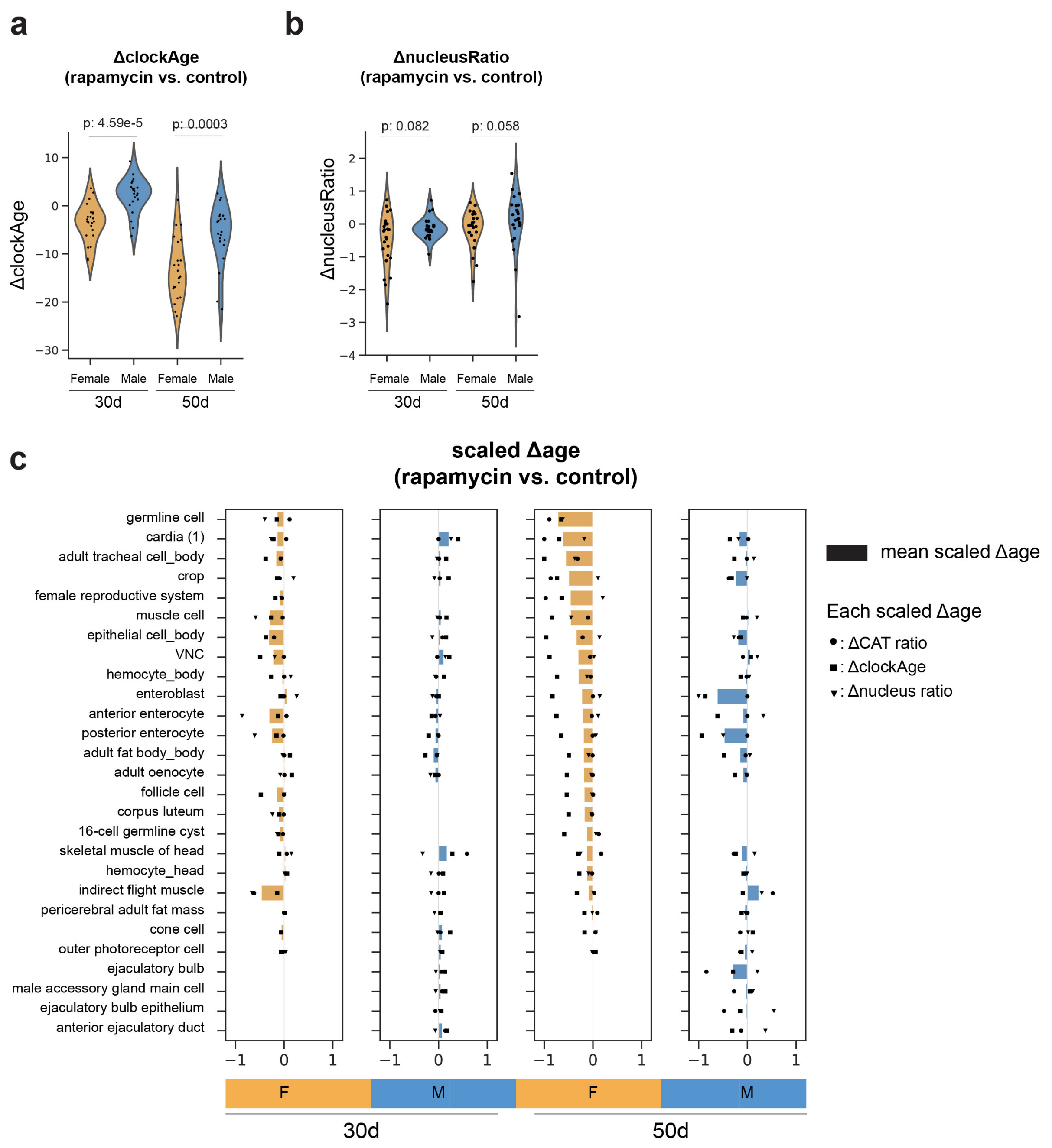
